# A mechanical atlas for Ascidian gastrulation

**DOI:** 10.1101/2022.11.05.515310

**Authors:** Siqi Liu, Patrick Lemaire, Edwin Munro, Madhav Mani

## Abstract

The intricate three-dimensional (3D) structures of multicellular organisms emerge through genetically encoded spatio-temporal patterns of mechanical stress. Cell atlases of gene expression during embryogenesis are now available for many organisms, but connecting these to the mechanical drivers of embryonic shape requires physical models of multicellular tissues that identify the relevant mechanical and geometric constraints, and an ability to measure mechanical stresses at single-cell resolution over time. Here we report significant steps towards *both* these goals. We describe a new mathematical theory for the mechanics of 3D multicellular aggregates involving the quasi-static balance of cellular pressures, surface tensions, and line tensions. Our theory yields a quantitatively accurate low-dimensional description for the time-varying geometric dynamics of 3D multicellular aggregates and, through the solution of a mechanical inverse problem, an image-based strategy for constructing spatio-temporal maps of the mechanical stresses driving morphogenesis in 3D. Using synthetic image data, we confirm the accuracy and robustness of our geometric and mechanical approaches. We then apply these approaches to segmented light sheet data, representing cellular membranes with isotropic resolution, to construct a 3D mechanical atlas for ascidian gastrulation. The atlas captures a surprisingly accurate low-dimensional description of ascidian gastrulation, revealing the adiabatic nature of the underlying mechanical dynamics. Mapping the inferred forces onto the invariant embryonic lineage reveals a rich correspondence between dynamically evolving cell states, patterns of cell division, and local regulation of cellular pressure and contractile stress. Thus, our mechanical atlas reveals a new view of ascidian gastrulation in which lineage-specific control over a complex heterogenous pattern of cellular pressure and contractile stress, integrated globally, governs the emergent dynamics of ascidian gastrulation.

## Introduction

A central challenge in biology is to understand how embryonic cells collectively shape tissues, organs, and whole embryos to determine organismal morphology. At the heart of morphogenesis are forces that are produced locally by individual cells, transmitted across cell-cell contacts, and resolved as global patterns of cell shape change, cell movement, and cell rearrangement. In recent decades, we have learned a great deal about the molecular machinery that cells use to produce and transmit forces within developing embryos [1, 2, 3]. We have also begun to learn how the deployment of this machinery is controlled in space and time by genetically programmed and self-organized patterns of gene expression and intercellular signaling [1, 2]. However, ultimately, the biochemical signals that “control” morphogenesis, and the molecular machinery they control, must speak the common language of mechanics; that is, they must act by controlling the local production and transmission of active forces, and the local material properties that determine the mechanical response of cells and tissues. In addition, there is increasing evidence that developmental and physiological control mechanisms are influenced by mechanical forces and deformations of cells and tissues, and transduce these into biochemical signals, to close mechanochemical feedback loops [4, 5, 6, 7, 8]. Thus the ability to measure mechanical forces over space and time in developing embryos is an essential prerequisite to understanding fundamental design principles for robust multicellular tissue morphogenesis.

In response to this challenge, a variety of experimental methods have been developed to measure forces in living embryos. One general class of methods involves local applications of force (e.g. through local mechanical indentation [9] or micropipette aspiration [10, 11] or optical or magnetic tweezers [12, 13]) or local disruption of mechanical continuity (e.g by laser ablation [14, 15]), coupled to the measurement of the resulting deformations. A complementary class of methods involves the observations of “force sensors” (e.g. molecular fret sensors, liquid droplets, or elastic beads) [16, 17], embedded within a force-producing tissue, whose deformations reflect internally generated forces. Importantly, all of these methods require the use of mathematical models to infer forces, and material properties, from observed displacements. These methods have provided valuable insights into the mechanics of cell and tissue morphogenesis at different spatial and temporal scales [18]. However, most are invasive and permit only sparse sampling at a few locations in space and time, limiting their use in mapping spatiotemporal patterns of forces that underlie the collective mechanics of multicellular tissue morphogenesis.

Recent advances in our abilities to image cell and tissue morphologies and deformations in living embryos have spurred the development of a third class of methods, commonly known as image-based force inference methods [19, 20, 21, 22, 23]. Image-based force inference methods use the shapes of cells in a multicellular aggregate themselves as sensors, using microscopic tissue-scale observations of cell shape and deformation to infer the underlying forces. Similar to other approaches, image-based force inference methods rely on physically-based mathematical models that relate the shapes of cells within tissue to the nature, organization, and magnitudes of the mechanical forces operating within individual cells and across cell boundaries. The central idea is to thus solve an inverse mechanics problem [20, 21, 22], to build a mapping from images of living tissues – the geometries of cells and cell-cell contacts – to the hidden variables such as tension along cell contacts and intracellular pressures, that define the mechanical state of a tissue. A key virtue of image-based approaches is that they are non-invasive, and by construction, offer the possibility of inferring forces at many simultaneously observed points within living embryos over time.

Thus far, with a few recent exceptions [24], methods for force inference have been implemented in two dimensions. Their applications have focused on the analysis of tissues, such as planar epithelia, in which it is assumed that the dominant mechanics operate in 2D, or which have sufficiently stereotyped geometries that a third dimension can be absorbed into a 2D description. More generally, however, the forces that regulate cell and tissue morphology are both patterned and resolved in three dimensions, and in these many cases, restriction to two dimensions fails to capture the relevant mechanical variables. For example, the physical conjugate to hydrostatic pressure – the cell volume – has no physically meaningful counterpart in two dimensions. Likewise, in most embryonic tissues, active contractile forces are organized with respect to surfaces and lines of contact (between 2 and 3 cells, respectively) which occupy and explore all possible orientations in three-dimension, defying a simple 2D representation. Recent developments attempt to address these challenges but do not account for the qualitative increase in mechanical and geometric complexity due to the fact that the active contractile forces of surfaces and lines of cell contact can be independently regulated.

Given these limitations of current force inference methods, and rapid advances in imaging approaches that allow the acquisition of high-resolution data on cell boundaries in three-dimensional over time in living embryos, the need to expand the capability for force inference to three-dimensional with sufficient mechanical complexity has become increasingly acute. However, doing so presents a number of serious theoretical and empirical challenges. First, it isn’t a priori apparent whether the inverse mechanical problem in three dimensions, inferring the mechanical state of the embryo from cellular geometries, is even a well-posed mathematical problem that permits inferring the mechanical unknowns from geometric knowns. Second, existing force inference methods rely on an image segmentation step in which an intermediate description of cell boundaries is extracted from the raw data. Unfortunately, the accuracy of the force inference is highly sensitive to the quality of the segmentation, and the segmentation becomes increasingly difficult in three-dimensional.

Here we describe a robust approach to three-dimensional force inference that overcomes these limitations. Our approach is based on a new general physical theory for the mechanics of close-packed three-dimensional cellular aggregates in which cell geometries are governed by the quasi-static mechanical balance of intracellular hydrostatic pressure forces, and contractile forces operating (differently) on curved surfaces and curved lines of contact between adjacent cells. We derive a mapping from the set of possible static geometries to values for pressure, surface tensions, and line tensions which is unique up to three scalar values or “zero modes”: Two of these define absolute scales for pressure and tension, respectively. A third quantifies the relative contributions of line tensions and surface tensions to defining a given geometry and highlights a form of morphogenetic flexibility in which different spatiotemporal patterns of force can be used to achieve identical outcomes. Applying our approach to in-silico-generated image data representing cellular aggregates at static mechanical equilibrium, we show that the global nature of the mapping and the underlying mechanics ensure a robust numerical solution, recovering the ground truth equilibrium geometry, and facilitating a robust inference of the underlying pressures and tensions. Finally, we apply our approach to light sheet data collected from membrane-labeled ascidian embryos during endoderm invagination, the first major step of ascidian gastrulation [25]. The remarkably good model fits these data strongly suggest that ascidian embryos operate close to mechanical equilibrium, i.e with adiabatic dynamics, revealing a surprisingly simple low-dimensional description of ascidian morphodynamics in terms of effective tensions and pressures - a three-dimensional mechanical atlas for gastrulation. In addition to confirming previous predictions about the forces that drive gastrulation, the mechanical atlas reveals surprising new insights into lineage-specific temporal control over intracellular pressure and the role of mechanical integration of pressure and tension across multiple tissues in driving ascidian gastrulation. Together our results establish a robust approach to three-dimensional force inference, grounded in a new three-dimensional mechanical theory of tissues operating close to mechanical equilibrium. They highlight the power of this tool to reveal fundamental physical design principles for morphogenesis, and its potential use to infer global mechanics in three dimensions in many other organismal and embryonic contexts.

## Theory

### Overview of Theory

To establish a foundation for three-dimensional force inference, we first introduce a mechanical theory for close-packed cellular aggregates in three-dimensional. Drawing upon a large body of experiments, and existing physical/mechanical theory, we assume that the shapes of cells within three-dimensional aggregates are determined by three biologically controlled mechanical inputs, which are schematized in Figure 1B. First, within each cell’s cytoplasm, an effective pressure resists changes in cell volume. While the dominant contribution to effective pressure is hydrostatic effects associated with an incompressible fluid-filled cytoplasm [26, 27], it may also include the isotropic effects of other active cellular processes, such as cytoskeletal assembly and motor activity [28]. Regardless of its manifold and complex molecular origins, or factors that control its magnitude, we consider the effective cellular pressure to be an isotropic stress that acts democratically to induce local curvatures at all the surfaces of the cell in question.

**Figure 1.**
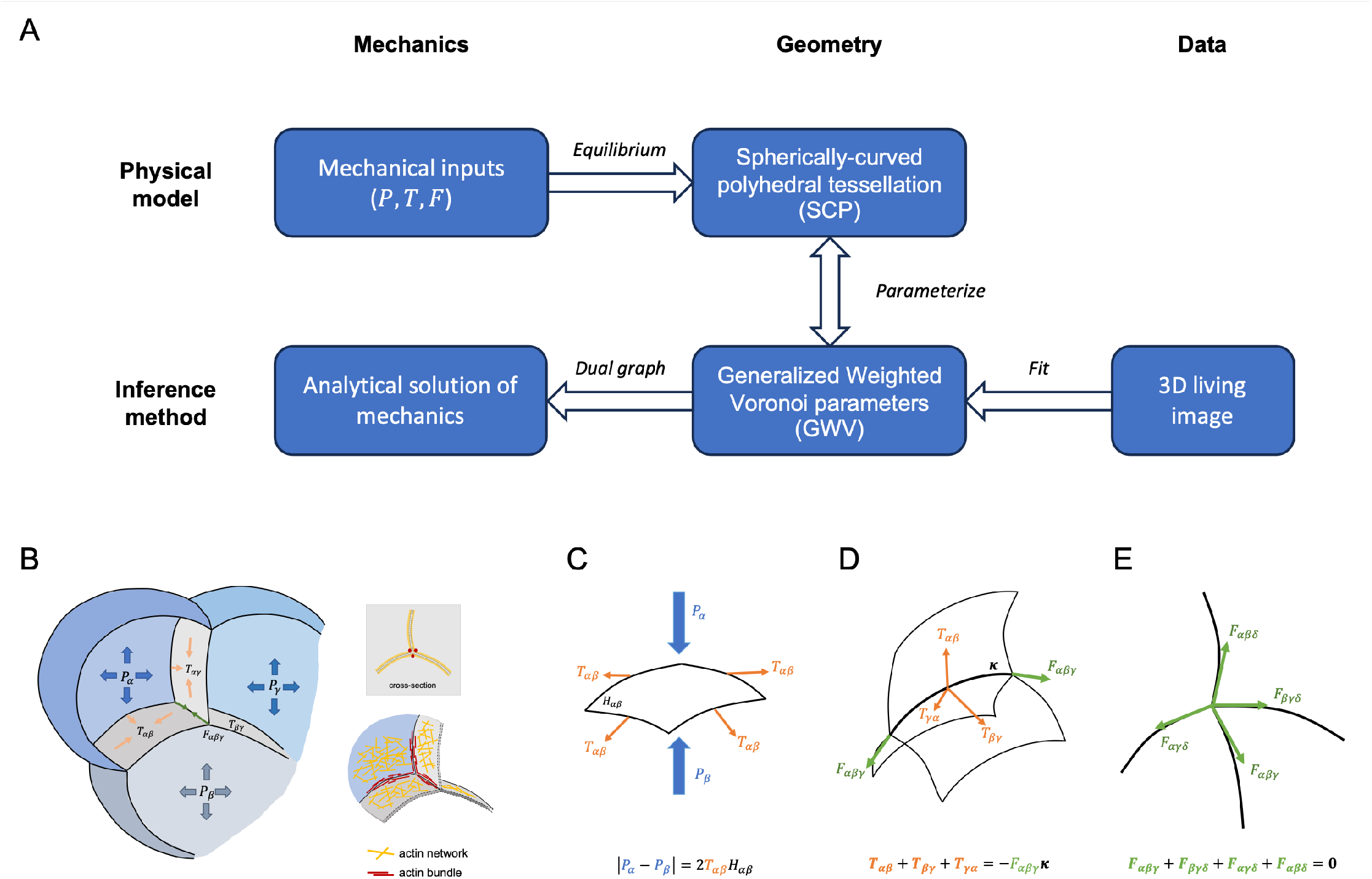
Mechanical theory for close-packed cellular aggregates in three-dimension. (A) A composite pipeline for the physical model and its associated force inference method. In the forward physical model, the values of mechanical inputs - pressures (*P*), surface tensions (*T*), and line tensions (*F*) - are given. Under the condition of static equilibrium (equations (1a)-(1c)), there exists a mapping from mechanics to geometry. The admissible geometries are spherically-curved polyhedral (SCP) tessellation, which can be parameterized by Generalized Weighted Voronoi (GWV) diagrams. Motivated by the physical model we can perform mechanical inference by numerically fitting the computational segmentation mesh obtained from live-imaging data to evaluate the best fit GWV parameters (see Figure 3). Following this parametrization of embryonic geometry we leverage our analytical results to evaluate the values of cell pressures, membrane, and line tensions (see Figure 2). (B)The mechanical factors accounted for are cell pressures (*P*, blue arrows), membrane surface tensions (*T*, orange arrows), and junction line tensions (*F*, green arrows) that have their origins in membrane-adjacent actin and adhesion networks and related structures (right panel). (C)The static equilibrium on a curved cell-cell contact with constant surface mean-curvature *H*_*αβ*_, according to force balance relations between isotropic surface tension (*T*_*αβ*_, orange arrows, tangent to the surface and orthogonal to the edge) and pressure difference (|*P*_*α*_*− P*_*β*_|, blue arrows, orthogonal to the surface) by Young-Laplace equation. (D)The static equilibrium on a curved junction with constant curvature *κ*_*αβγ*_, according to force balance relations between a homogeneous line tension (*F*_*αβγ*_, green arrows, tangent to the junction) and three isotropic surface tension (*T*_*αβ*_, *T*_*βγ*_, *T*_*γα*_, orange arrows) by 1d version of Young-Laplace equation. (E)The static equilibrium on a vertex, according to force balance relations between four line tensions (*F*_*αβγ*_, *F*_*βγδ*_, *F*_*αγδ*_, *F*_*αβδ*_, green arrows).

Second, effective surface tensions act along free cell surfaces and cell-cell contacts to oppose the curvature induced by effective cellular pressures. A large body of work suggests that the key determinants of surface tension in close-packed tissues are contractile forces produced by the cortical actomyosin cytoskeleton, working against a passive resistance of the cortex to deformation [1, 29]. Finally, surface tensions act to induce curvature along junctions where two or more cells meet, and line tensions within those junctions oppose this curvature. Again, the key determinant of line tensions along cell contacts is thought to be contractile forces produced by actomyosin networks that assemble adjacent to cell-cell contacts. In contrast to simple soap bubble foams where surface tension and line tension are co-determined, actomyosin contractility can be independently controlled on cell contacts and junctions in multicellular aggregates [30], including ascidian embryos [31, 32, 33]. Thus here we account for surface tensions and line tensions as separate quantities, admitting the necessary physical and mathematical complexity.

To make a force inference tractable, we make several additional empirically motivated assumptions. First, measurements of cell or tissue response to laser ablation in a variety of different tissues are rapid (tens of seconds or less) [34], compared to the timescale of morphogenesis (minutes to hours), suggesting that the majority of mechanical forces in play are in static equilibrium with each other on these longer timescales – i.e. the dynamics are adiabatic. Second, while active forces probe the elastic responses of the cytoskeleton at short timescales, these forces are dissipated by cytoskeletal remodeling and turnover at longer timescales. These remodeling and turnover dynamics motivate posing a fluid constitutive model, in particular the Young-Laplace relation, that stitches together differential cell pressures, surface tensions, and line tensions in a static multicellular aggregate. Third, while there may be spatial and/or temporal heterogeneities and anisotropies within a single cell-cell contact or junction, we will assume that these equilibrate, or “average out” at longer times. Similarly, we will assume that local variations in pressure within a single cell’s poroelastic cytoplasm [35] equilibrate or average out at longer times. Thus, on morphogenetic timescales, we will assume: (a) that the cell surfaces and lines of contact behave effectively like simple fluids; (b) that single, time-dependent, numbers quantify the effective pressure within each cell, the surface tension along each cell contact, and the line tensions along each junction; and (c) that at any moment in time, the system operates close to a static mechanical equilibrium.

With these assumptions, the physical rules that govern cell shapes can mathematized as follows: On each free cell surface or cell-cell contact, the balance of effective pressures and surface tension can be described by the well-known Young-Laplace (YL) relation. Intuitively, the YL relation states that an effective pressure difference across a fluid surface produces a force that acts to curve that surface; surface tension produces an opposing force that acts to straighten the surface; and these forces are balanced when the mean curvature of the surface equals the ratio of the effective pressure difference and the effective surface tension. Using indices *α, β, γ* to label individual cells, the balance of stresses normal to surfaces (Figure 1C) is given by the Young-Laplace equation:

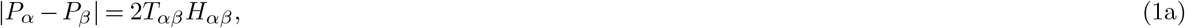

where *P*_*α*_, *P*_*β*_ are effective pressures, *T*_*αβ*_ is surface tension, and *H*_*αβ*_ is the mean curvature of membrane surface. Importantly, the assumption that surface tension is homogeneous and isotropic within a given surface implies that only surfaces of contact with a constant mean curvature (CMC) [36] are permitted by this mechanical model. The potential class of surfaces with CMC is vast and complex. Qualitatively, the membrane geometries observed in embryos are simple compared to this library of potential CMC surfaces, and hence going forward we limit our attention to the spheres, which are the simplest class of CMC surfaces.

In a physically and mathematically analogous way, surface tensions on cell contacts (or free cell surfaces) produce forces that act to curve junctions where three surfaces meet. These in turn are opposed by line tensions along the junction. An analogous YL relation states that the curvature along each junction is the ratio of the effective surface and line tensions. Thus, the junctional balance of force (Figure 1D) is described by a one-dimensional Young-Laplace equation:

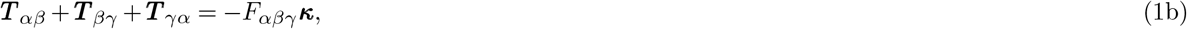

where ***T*** _*αβ*_ is a force acting normal to the junction whose magnitude is the surface tension *T*_*αβ*_, *F*_*αβγ*_ is the junction’s line tension, ***κ*** is the *curvature vector* of the junction. Again, the assumption that line tension is homogeneous along a given junction implies that only junctions of constant curvature, i.e. sections of circles, are permissible. With this constraint, the curvature vector *κ* is uniquely defined as the vector of constant length lying normal to the junction and pointing towards the center of the circle.

Finally, the line tensions along each junction produce forces at vertices where four junctions meet. The assumption that all mechanical forces are in static balance with one another implies that these forces must sum to zero (Figure 1E):

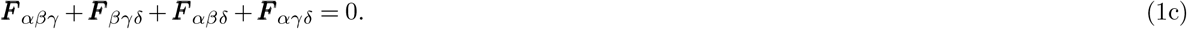

Equations (1a)-(1c) constitute a mechanical theory for compact three-dimensional cellular aggregates at quasi-static mechanical equilibrium. In particular, they define relationships between mechanics and cell geometry in static mechanical equilibrium, given the underlying assumptions of our theory. In accordance with the assumption that surface and line tensions are homogeneous and isotropic across individual contact surfaces or junctions, the only shapes we allow at static equilibrium are those that belong to the special class of Spherically Curved Polyhedral tessellations (SCPs), in which free cell surfaces or cell-cell contacts are sections of spheres, and junctions are sections of circles (Figure 2). This reduction in complexity facilitates our approach to force inference in two critical ways: First, as we will show below, it allows us to identify a set of independent descriptors of geometry, and analytical formulae, from which the mechanical parameters can be computed directly up to three undetermined constants. Second, and equally important, it supplies an intermediate description of cell geometry onto which real data can be mapped, and it does so in a way that allows robust quantitative assessment of the applicability or fit, of the theory to data from a particular system, and thus the validity of the force inference approach for that system.

**Figure 2.**
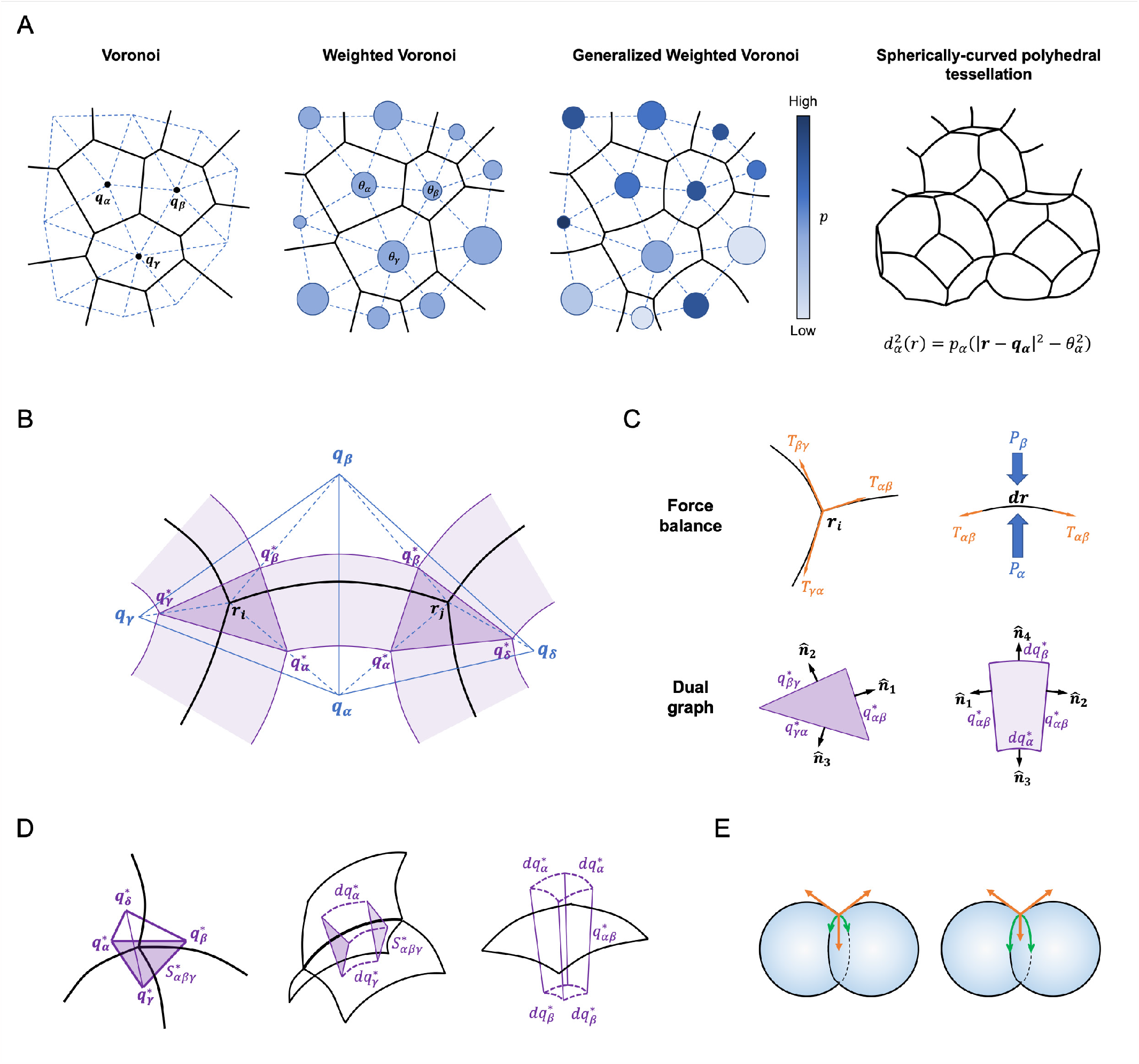
Inferring mechanics from geometry. (A)Parametrization of Spherically-curved polyhedral tessellation by Generalized weighted Voronoi diagram, visualized in 2D. In the first panel, given the Voronoi site (***q***, black dots) for each cell, the Voronoi diagram is constructed according to Euclidean distances 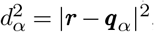 where cell shapes are polygons. In the second panel, given the additional weights (*θ*, the radius of the blue circles) for each cell, the weighted Voronoi diagram is constructed according to the distance 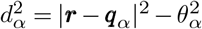, where cell shapes remain polygons but their sizes vary. In the third panel, given the power (*p*, light to dark blue color of circles) for each cell, the Generalized Weighted Voronoi diagram is constructed according to 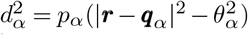, where cell boundaries can now adopt the shape of circular arcs. (See SI Part I.D for details.) In the fourth panel, we illustrate this construction in 3-dimensions. (B)Two-dimensional Circular Arced Polygon (CAP) tessellation and the construction of its dual graph. The two ends of a dual line (***q***^*\**^, purple points) for a general point ***r*** on cell boundary is defined by 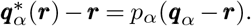. On a vertex (***r***_*i*_), three dual points form a dual triangle (dark purple). Along a cell membrane, the dual line sweeps out a curved quadrilateral (light purple). (C)(D)(E) Inferring mechanics from geometry. (C)Two-dimensional force balance relations and the corresponding dual graphs. At a vertex (***r***_*i*_, top-left), three surface tensions (*T*_*αβ*_, *T*_*βγ*_, *T*_*γα*_, orange arrows, tangent to the surface) are under static equilibrium, according to the force balance relation ***T*** _*αβ*_ + ***T***_*βγ*_ + ***T***_*γα*_ = **0**. This static equilibrium of forces corresponds to the divergence theorem of the corresponding dual triangle (dark purple, bottom-left) 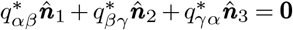. Along a surface element (*dr*, top-right), the surface tension (*T*_*αβ*_, orange arrows) and pressures (*P*_*α*_, *P*_*β*_, blue arrows, orthogonal to *dr*) are under static equilibrium, according to Young-Laplace relation 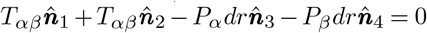. This balance of forces corresponds to the divergence theorem of the appropriate dual quadrilateral (light purple, bottom-right) 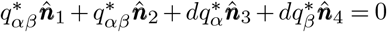. (D)Three-dimensional dual graphs. The force balance relation of four line tensions on a vertex in Figure 1E corresponds to the divergence theorem of the dual tetrahedron (left panel). The force balance relation between line tension and three surface tensions along a junction in Figure 1D corresponds to the divergence theorem of the dual triangular tube (middle panel). The force balance relation between surface tension and pressure difference on a surface element in Figure 1C corresponds to the divergence theorem of the dual curved shell (right panel). See SI Part II.C for details. (E)A two-cell example of the different force configuration on the same geometry under the third mechanical zero mode. In the left panel, a small value for the line tension on the middle junction (green arrow) and a large surface tension on the middle cell-cell contact (orange arrow) are under static equilibrium. In the right panel, a large value of line tension on the middle junction (green arrow) and small surface tension on the middle cell-cell contact (orange arrow) are also under static equilibrium, resulting in the same geometry.

## Inferring Mechanics from Geometry

### Overview

Equations (1a)-(1c) define relationships between mechanics and cell geometry that must hold at mechanical equilibrium, given the underlying assumptions of our theory. The fundamental question we seek to address is: given an estimate of the geometry of all cells in an embryo obtained from three-dimensional imaging, can we infer its mechanical parameters? Stated as an inverse mechanics problem: Given Equations (1a)-(1c) and imaging data, can we infer values of effective pressures, surface tensions, and line tensions for all individual cells, surfaces, and junctions?

At first glance, the solution to this inverse problem seems trivial since equations (1a)-(1c) are linear! However, the notation in these equations hides the fact that the system of equations is highly coupled across neighboring vertices, lines, surfaces, and cells. Additionally, since the only equilibrium geometries we consider here are SCP tessellations, the positions of vertices and the curvatures of surfaces and junctions are not independent. For example, the curvature of a junction and the positions of its two vertices are not independent since they must define a section of a circle. Similarly, the junctions that circumscribe the boundary of a surface and its curvature cannot be independent since they must collectively reside on a section of a sphere. Thus, *a priori*, it is not actually clear whether the highly coupled systems of equations are solvable – i.e. whether there are enough constraints to determine all of the unknown mechanical parameters.

As we outline below and derive more fully in the SI Part I, the key to solving the inverse mechanics problem lies in identifying the geometric constraints imposed by an SCP tessellation, and using these to reparameterize the description of geometry in terms of independent variables. We show that this can be done by equating the family of all SCP tessellations to a class of Voronoi diagrams known as generalized weighted Voronoi (GWV) diagrams. This yields a path to solving the inverse problem in two steps. First, using a parameterization given by the GWV construction, we determine the SCP tessellation that most closely fits the observed geometry. Importantly, the goodness of this fit constitutes a test of our mechanical assumptions. Second, using the GWV construction, we show that equations (1a)-(1c)) can be solved analytically to obtain pressures and tensions from the geometric parameters that describe an SCP tessellation. These solutions are not unique, but instead furnish values for the mechanical parameters up to three global undetermined constants, which we refer to as zero modes. As we detail further below, one corresponds to the background pressure which can be set to zero without loss of generality. The second corresponds to the absolute magnitude of physical forces, which cannot be inferred from geometry alone, but which could be measured for a given system. The third reflects the relative contributions of surface and line tensions to producing a given equilibrium geometry. As such, it reveals a form of biological redundancy that embryos may exploit through differential control of junctional or cortical contractility to achieve the same morphogenetic outcome in different ways.

### Generalized weighted Voronoi constructions parameterize SCP tessellations

SCP tessellations are comprised of vertices connected by junctions that are sections of circles, which are themselves connected by curved cell surfaces that are sections of spheres. The curvature of a given junction is codetermined by the curvatures of the surfaces that it intersects. We can associate with each curved surface a unique point - its centroid - that lies at the center of the sphere that contains that surface. Therefore, an SCP tessellation is described by the positions of all of its vertices and centroids. However, these positions are interdependent. The vertices of a given membrane must be equidistant from its centroid. As for polyhedral tessellations with planar membranes, the positions of all vertices that intersect a given surface must lie on a plane. Finally, as we show in the SI (Part I.C), the centroids of three adjacent membranes must be co-linear. Together, these constraints imply that for SCP tessellations, the number of degrees of freedom, i.e. the number of independent quantities required to specify the geometry for a specific SCP tessellation is much less than the number of vertex and centroid positions. In fact, as we show in the SI (Part I.C), if *N*_*C*_ is the number of cells in an aggregate, then an SCP tessellation has 5*N*_*C*_ independent degrees of freedom.

As we show formally in SI (Part I.D), an alternative way to describe an SCP tessellation is to generalize the classical method for constructing Voronoi polygons [22]. Given a set of *N*_*C*_ points *q*_*α*_ in three dimensions, we identify a region surrounding each point as the set of points ***r*** lying closer to that point than any other, where distance is defined as a classical euclidean distance 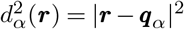. This defines a set of polyhedral regions whose shared boundaries lie equidistant between their center points (Figure 2A, first panel). Modifying the distance function by adding a squared weight 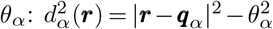introduces tunable variation in the sizes of the polyhedrons (Figure 2A, second panel). Finally, multiplying the weighted distance function by a factor 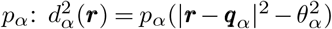 induces tunable curvature of polyhedral boundaries (Figure 2A, third panel).

It follows that, for each polyhedral cell, there are exactly 5 independent quantities - the three coordinates of the center point *q*_*α*_, and the values of weights *θ*_*α*_ and powers *p*_*α*_. Thus an SCP tessellation of *N*_*C*_ cells maps to a unique (see SI Part I.D for the formal argument) generalized weighted Voronoi (GWV) diagram (Figure 2A, fourth panel), which is defined by 5*N*_*C*_ independent parameters Ψ = 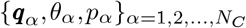. Henceforth, we will omit reference to the GWV construction and use Ψ to represent the set of geometric parameters that uniquely describe a given SCP tessellation.

### An analytical solution to the inverse mechanical problem

To solve the inverse mechanical problem, we require an analytical solution of the equations of motion, (1a)-(1c), that relates the parameters of a GWV construction to the effective pressures and tensions that produce a given geometry. We derive the solution for the three-dimensional problem in the SI Part II. Here, for the sake of pedagogy, we explain the solution for the simpler, and easier-to-visualize, scenario of two dimensions. The approach we use to derive the two-dimensional solution is distinct from that taken by Noll et al [22], and generalizes readily to three dimensions. In the two-dimensional scenario, the system’s mechanical inputs are pressures and membrane tensions, alone. The mechanical balance at vertices and membranes can be described by ***T*** _*αβ*_ + ***T***_*βγ*_ + ***T***_*γα*_ = **0** and |*P*_*α*_ − *P*_*β*_|= *T*_*αβ*_*/R*_*αβ*_, respectively. Here, *R*_*αβ*_ refers to the radius of the membrane edge. For a two-dimensional system with cell pressures and membrane tensions, the static equilibria correspond to Circular Arced Polygon (CAP) tessellations, which are the two-dimensional analogs of SCPs. CAP tessellations are shown in black in Figure 2B and can be parameterized by a two-dimensional GWV tessellation, with 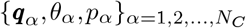 as the parameters.

In this two-dimensional context we now describe the construction of the most central mathematical objects in our analytical solution – the dual graph – that will form the basis for the bridge between the parameters of the SCP tessellation and the mechanical forces. Following the schematic shown in Figure 2B, for any point ***r*** on cell-cell boundary - membrane or vertex - of cell *α*, we define the dual point as 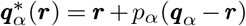. A corresponding dual point exists in the adjacent cell, 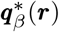. The two dual points corresponding to the point ***r*** on the membrane form the dual line 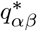 corresponding to the membrane between cell *α* and *β*. It is easy to show in the SI (Part II.B) that the dual line is perpendicular to the membrane tangential direction at ***r***, and the length of the dual line is constant along the membrane. As shown in purple in Figure 2B, at a vertex, three dual lines form a dual triangle. Furthermore, along the membrane, the dual lines sweep out a curved quadrilateral.

The analysis of the dual graph that follows relies heavily on several elementary geometric facts, one of which is a special case of the divergence theorem that is derived in SI (Part II.A). The important implication of this is that the integral of the surface normal over the boundary of a closed region is zero: 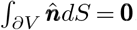. In the two-dimensional case, the integral of the normal over a closed shape is zero: 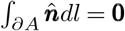. We are now positioned to apply the divergence theorem to the dual construction. To help follow the mathematical procedure we have schematically excised the triangle and quadrilateral and displayed them in Figure 2C, along with the force balance relations they correspond to. Applying the two-dimensional divergence theorem to a dual triangle, gives us 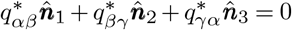. We notice that the three vectors have same directions as the corresponding membrane tension balance 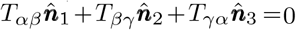Therefore, the length of the dual line is proportional to the membrane tension: 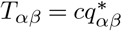, where *c* is a single constant for all membranes in a tessellation. Now considering the edge element *d****r*** along the membrane adjacent to cell’s *α & β*, the dual shape is a curved quadrilateral as shown in Figure 2C. Applying again the two-dimensional divergence theorem to the quadrilateral, we obtain 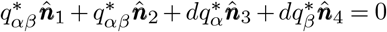. Observing that the vectors have same directions as the corresponding tension-pressure balance on *dr*, 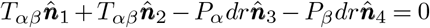, which is the differential statement of the Young-Laplace equation. Since 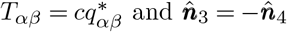, the pressure difference can be rewritten as 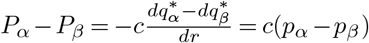. Finally giving us the pressure in cell *α, P*_*α*_ = *cp*_*α*_ + *b*, where *c* is the same constant as before, and *b* is a single global constant setting the overall baseline of pressures.

The machinery described above extends naturally to three-dimensional situations, as detailed in the SI (Part II.C) and illustrated in Figure 2D. In three dimensions, the dual point of a point ***r*** on the boundary is still given by 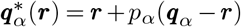. As a result, the closed dual shape corresponding to a vertex takes the form of a tetrahedron, while the closed dual shape corresponding to an edge is a triangular tube and the dual shape corresponding to a membrane is a shell with a certain thickness. Applying the three-dimensional divergence theorem to these shapes and relating them to the mechanical inputs, we determine that the line tensions (*F*_*αβγ*_) are proportional to the areas of the triangles, the membrane tensions on an edge element (*T*_*αβ*_*dl*) are proportional to the tube’s lateral faces, and the cell pressures on a membrane element (*P*_*α*_*dS*) are proportional to the shell’s inner or outer surface. A detailed derivation appears in the SI (Part II.C). Here we present the final analytical solutions:

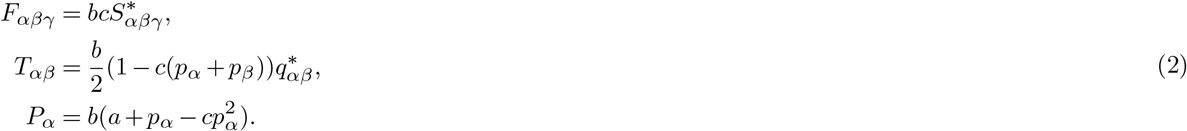

These three equations map the parameters that describe an SCP tessellation to the set of pressures, surface tensions, and line tensions that make that SCP tessellation a mechanical equilibrium, a map that is unique up to the values of three global unknown constants denoted as *a, b* and *c*. These constants cannot be determined by geometry alone and thus are unknown constants that can change in time. Crucially, they are uniform in space. It is this central result that makes it possible to infer a mechanical atlas of forces from the description of an embryo’s geometry. We highlight that the map is nonlinear and in no way could one intuitively derive it based on simplistic mathematical and physical considerations.

### Stress tensor

The forces inside each cell and on its boundaries - membranes and junctions - together provide stresses on the cell. Inside a cell, the cell volume is under an isotropic stress, pressure, ***σ***_*P*_ = *P*_*α*_ ***I*** _3_ (where ***I*** _3_ denotes the identity matrix in three-dimensions). On a membrane, surface tension generates an in-plane mechanical stress ***σ***_*T*_ = *T*_*αβ*_ ***I***_2_ (where ***I***_2_ is the in-plane projection of identity matrix). Along each edge, line tensions generates a tangential stress ***σ***_*F*_ = *F*_*αβγ*_ ***I***_1_ (where ***I***_1_ is one-dimensional projection of identity matrix). Accounting for all of these, the cellular stress tensor is defined as

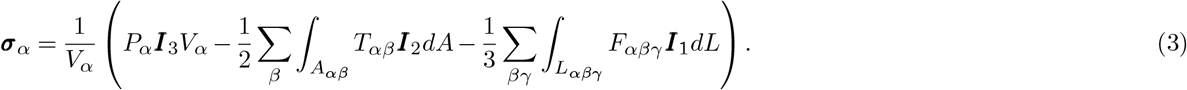

Here the 1*/*2 and 1*/*3 factors represent the share of the stress on boundaries by two or three cells. The minus signs before the second and third terms denote that the surface stress and line stress both contribute contractility of ***σ***_*α*_, which can be separately expressed as the cellular contractile stress tensor:

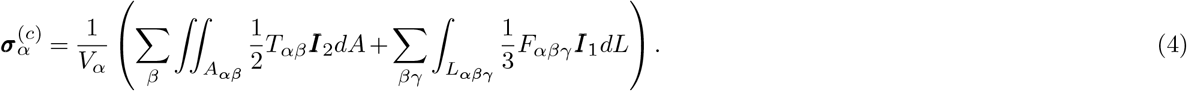

By construction, the cellular stress tensor is a 3*×*3 symmetric tensor, and the scaling by the volume of the cell ensures that the tensor itself is an intensive quantity. This cell-centric mechanical stress tensor holds coarse-grained information regarding its patterns of mechanical forces. Although the individual forces are modeled to be isotropic, the cellular stress tensor has both isotropic and anisotropic components owing to the geometric considerations. While the isotropic part is straightforwardly quantified by the trace of the matrix, the traceless anisotropic part can be quantified by the scalar shear stress termed von Mises Stress, 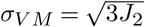, through the second characteristic *J*_2_ of the deviatoric matrix.

### Zero modes

The constants *a, b*, and *c* represent three undefined scales in the expressions of forces that produce a given SCP tessellation. In physical terms, there are three zero modes in the correspondence between the mechanical degrees of freedom identified by our physical model and the geometric degrees of freedom that describe all possible SCP tessellations. Stated differently, a single SCP tessellation corresponds to a three-continuous-parameter family of mechanical states. The scale *a* determines the ambient pressure scale, which can be set to 0 without loss of generality. Physically speaking, this freedom reflects the fact that only differences in pressures enter our model, but not the absolute scale of pressures themselves. The scale *b* can be understood as a factor that determines the overall scale of forces in physical units. An inference based purely on geometry cannot inform us of the units in which the forces are expressed. To summarize, two of the three zero modes deal with the ambient scale of pressure and the units of force [22]. These constants can change in time but are uniform across an embryo at a given time point.

The third zero mode is parameterized by *c* and reflects a novel consequence of the three-dimensional interplay between surface and line tensions. The physical nature of this zero mode can be seen by considering a simple two-cell system as shown in Figure 2E. Here the two cells have equal pressure (ensuring that the contact between the two cells is flat), and the angles between the three membranes are equal at the junction. This geometry can be supported by a continuous one-parameter family of mechanical states where the line tension of the central contact, and its membrane tension, can covary with each other in a negative manner. Focusing on a single location along the line of contact, both the membrane and line tension of the central membrane provide a force towards its center. Thus, the same arrangement of angles can be sustained by a weaker/stronger line/membrane tension, or vice versa.

Generalizing (see SI Part II.D for details), any given SCP tessellation can be sustained by a one-parameter family of mechanical states, that places more or less relative weight on the contributions of surface and line tensions. The value of the c constant can change in time, but generically adopts a fixed value in space – except for a special case (see SI Part II.D). This isn’t a result that can be gleaned from simplistic mechanical or physical arguments and is a novel feature of three-dimensional aggregates with independent line and membrane tensions that is revealed by the mathematics presented here. Going further, assuming that all surface and line tensions in the system are generated by contractile systems (e.g the actomyosin cytoskeleton [1]), that do not generate extensile forces, provides bounds on the values that *c* can take on, 0 ≤ *c* ≤ *c*_*max*_, where *c*_*max*_ = 1*/* max_*α,β*_ (*p*_*α*_ + *p*_*β*_).

From the practical standpoint of force inference, determining the value of *c* requires information beyond the purely geometric description furnished by light-sheet imaging data. This information could be obtained by additional measurement, but here we are limited to inferring the mechanical stresses in an embryo up to a single, potentially time-varying, global constant.

## Fitting the data

### Overview

Given the assumptions of our theory for three-dimensional aggregates, the above relations provide the map from parameters Ψ ={***q***_*α*_, *θ*_*α*_, *p*_*α*_ }_*α*=1,2,…,*C*_ that describe an SCP tessellation to its mechanical state - up to zero modes that cannot be identified by any image-based approach. To infer the mechanical stresses underlying an observed embryo’s geometry, we must first map the observed geometry to an SCP tessellation. Of course, there is no guarantee that real embryos satisfy the assumptions of our theory. Indeed, the extent to which an observed three-dimensional geometry is well-approximated by the much simpler geometry of an SCP tessellation provides a test of whether the theory’s assumptions hold for a given embryo. Below, we outline methods (see SI Part III for details) to globally project three-dimensional image data representing cell boundaries within an embryo onto the closest SCP tessellation and to quantify the degree of mismatch between the observed and projected geometries.

### A minimization statement

Quantitatively speaking, the segmented image obtained from live embryo imaging provides the coordinates of membrane pixels, ***r***_*i*_, from which we seek to reconstruct a simpler geometry - an SCP tessellation. Denoting the centroid and radius of membrane sphere by ***ρ***_*αβ*_ and *R*_*αβ*_, the deviation of the pixel from sphere is *ϵ*_*i*_ =|***r***_***i***_−***ρ***_*αβ*_ |−*R*_*αβ*_ (Figure 3A). Since ***ρ***_*αβ*_ and *R*_*αβ*_ are both functions of the geometric parameters Ψ = {***q***_*α*_, *θ*_*α*_, *p*_*α*_*}*_*α*=1,2,…,*C*_, that define an SCP tessellation, we can recover the geometric parameters by least-squares fitting. Specifically, we minimize the mean-squared-deviation (MSD) function,

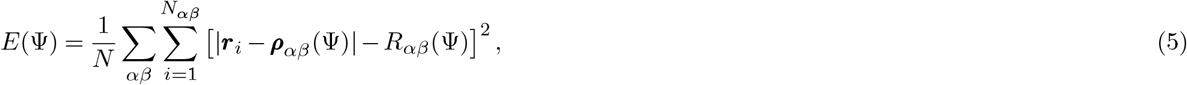

where *N* = ∑_*αβ*_*N*_*αβ*_ denotes the total number of pixels, to find the ‘closest’ SCP tessellation to the observed data. Our minimization statement falls into the class of nonlinear optimization problems, requiring an educated initial guess for the desired parameters that takes advantage of the properties of SCP tessellations – see SI (Part III.A) for further details. The minimized value of *E*(Ψ) is a global measure of the average deviation of the empirically observed geometry and the best-fit SCP tessellation. However, as we will demonstrate below, our approach provides finer-grained spatial information pertaining to the errors in our approximation.

**Figure 3.**
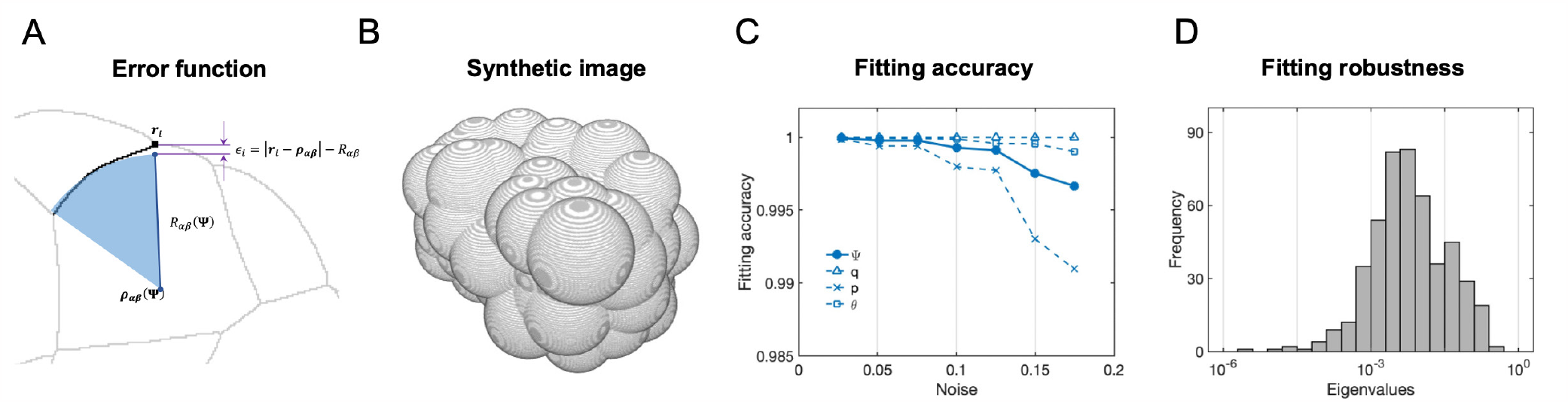
Method, precision, and robustness of image fitting for geometric parameters. (A) The definition of the error function in a 3-dimensional image labeling membrane pixels (in a 2-dimensional cross-section). For given geometric parameters **Ψ** of an SCP tessellation, each membrane shape is a section of a sphere (blue, centroid ***ρ***_*αβ*_ (**Ψ**), radius *R*_*αβ*_(**Ψ**)). For each pixel on the membrane (black square dot, ***r***_*i*_), the deviation (purple,ϵ_*i*_) to the sphere is defined by ϵ_*i*_ = |*r*_*i*_*−ρ*_*αβ*_|*− R*_*αβ*_. (B)A synthetic 3-dimensional image of cellular aggregate with know parameters Ψ^*\**^ and Gaussian noise on pixels (with standard deviation *δ*). (C)Accuracy of fitted parameters Ψ, relative to ground truth, Ψ^*\**^ on synthetic images with different levels of Gaussian noise (*δ*). Broken lines show the accuracy of the fitted three categories of Voronoi parameters (site ***q***, weight *θ*, power *p*). (D)A sensitivity analysis on synthetic data showing the fitting roboustness. At the minimum of the error function, we evaluate the eigenvalues corresponding to the linear response to noisy perturbations of pixels. All the eigenvalues are less than 1, indicating the robustness of the numerical scheme to noise.

### Precision and robustness of data fitting

Before applying our approach to a specific biological case, we first sought to assess the robustness and precision of the data-fitting scheme when applied to a comparably complex scenario *in silico*, where we have direct access to the underlying ground-truth. Given random parameters Ψ_0_, defining a specific SCP tesselation for *C* cells, we can identify the locations of all membrane pixels to produce a three-dimensional image. After introducing Gaussian noise into all pixel locations, we generate synthetic noisy images of cellular aggregates, as shown in Figure 3B. Our fitting scheme recovers a best-fit guess Ψ that we can compare with Ψ_0_, the ground truth. The results of this comparison (Figure 3C) report the degree of mismatch, or error, between the parameters of the SCP tesselation inferred by our scheme, relative to ground truth, as a function of the degree of noise injected into the geometry. Although the inferred parameters of the SCP tesselation display varying degrees of susceptibility to noise, the fitting scheme recovers parameters with up to 99% accuracy with as much as 20% noise injected into the location of membrane pixels. See SI (Part III.B) for more details.

To further assess the robustness of our numerical fitting scheme, we perform a sensitivity analysis, asking to what extent the inferred parameters of an SCP tessellation vary when we impose small perturbations of the data around some reference configuration. To assess this mathematically, we analyze the eigenvalue distribution of the system’s linear response [20]. An inference scheme is robust when all its eigenvalues are smaller in magnitude than 1, ensuring that no perturbations to the system can generate disproportionately large deviations in the values of inferred parameters. In particular, at the minimum of the MSD function, *δE*(Ψ, ***r***) = 0, thus, for any ***r***_*i*_, there must be *δϵ*_*i*_ = *δ*[|***r***_*i*_ −***ρ***_*αβ*_ (Ψ)| −*R*_*αβ*_(Ψ)] = 0. This permits us to derive a local linear approximation, *Kδ****r*** + *Mδ*Ψ = 0, where *K* = *∂* ***ϵ*** */∂* ***r*** and *M* = ***∂*** *ϵ /∂*Ψ are two matrices. As such, the relation between observed pixels perturbation *δ****r*** and geometric parameter deviation *δ*Ψ is given by 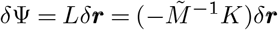, where 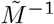 is the pseudoinverse matrix of *M*. For synthetic data, we compute the eigenvalues *λ*_*i*_ of *L* and find they are all less than 1 in magnitude (Figure 3D). This indicates that the fitting scheme is robust to noise.

## Results

We then turned to ascidian embryos (Figure 4) as a powerful in vivo test case for our three-dimensional force inference methods. Early ascidian development is characterized by the early specification of major embryonic axes, an invariant lineage (Figure 4B), and a highly stereotyped sequence of morphological changes governed by the timing and orientation of cell divisions and by lineage-specific changes in cell shape [37]. Recently, Guignard et al combined multi-view light-sheet imaging and image segmentation to reconstruct the boundaries of all cells within the ascidian embryo at 2-minute intervals during early ascidian development with isotropic resolution [25]. Here, we focus on the first part of gastrulation in which the ten endodermal precursor cells (yellow in Figure 4A&B) invaginate to form a large indentation on the vegetal side of the embryo, accompanied by a large-scale deformation of surrounding mesodermal and ectodermal tissues, encompassing the entire embryo. We applied our three-dimensional force inference to the segmented light sheet data from Guignard et al to produce a mechanical atlas for early gastrulation, mapping spatiotemporal patterns of force production onto the embryonic lineage to identify both lineage-specific and collective (embryo-wide) modes of morphogenetic control.

**Figure 4.**
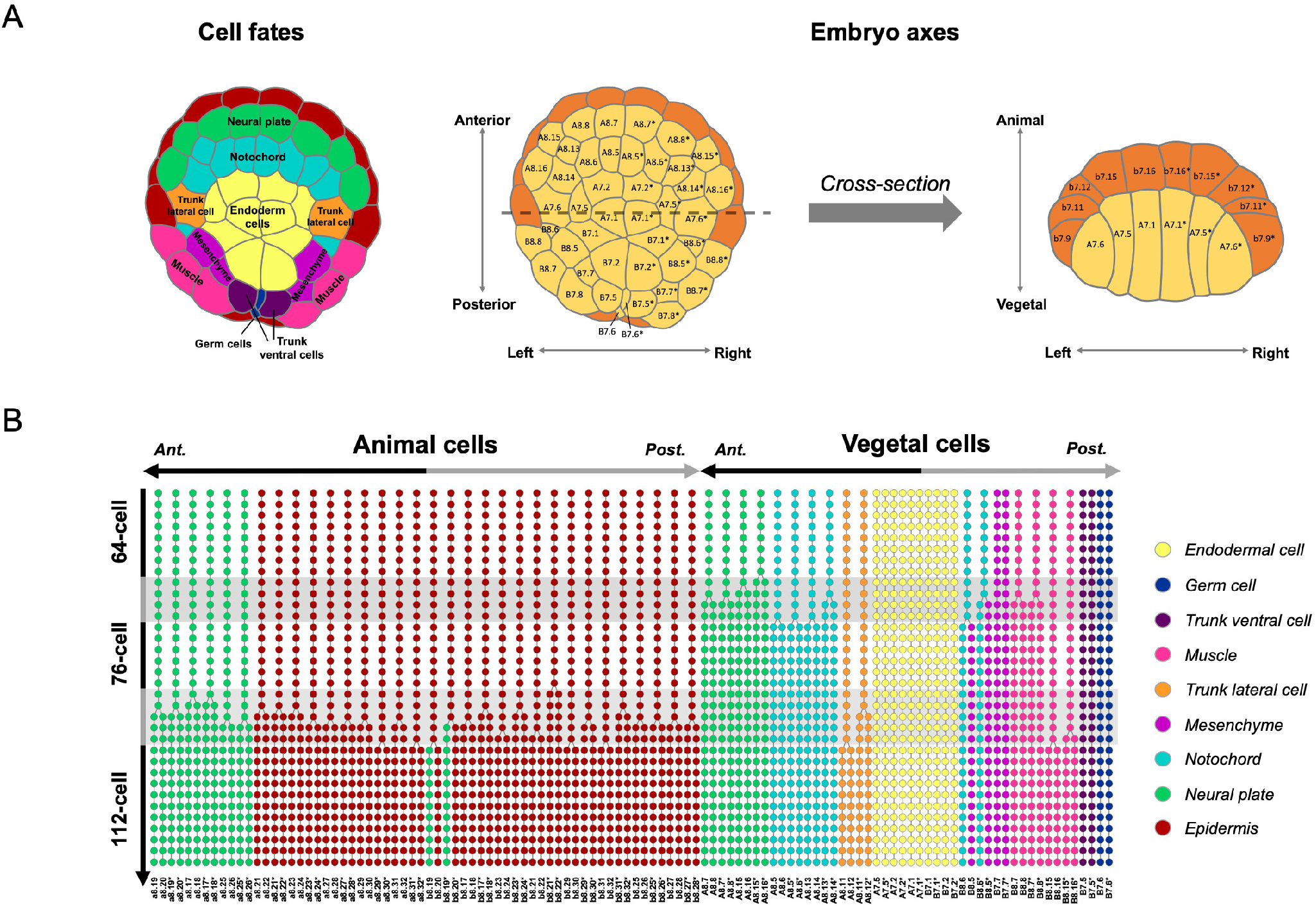
Early gastrulation in the *Phallusia mammillata* embryo. (A) Cell fates and embryonic axes of the embryo at the 76-cell stage. In the first panel, cell fate is indicated by colors. The second panel indicates the names of vegetal cells (yellow) and their organization along the anterior-posterior and left-right axes of the embryo. The third panel presents a cross-sectional view of the embryo along the broken line in the second panel, indicating the names of cells of the vegetal and animal layers (orange). (B) A lineage map of cell fates through the process of gastrulation in the *Phallusia mammillata* embryo from [25]. Each dot in the lineage represents a cell in time between the 64-cell and 112-cell stages, with the color indicating its fate. Timepoints are separated by 2 minutes. Cell names are indicated at the bottom at the 112-cell stage. Cells are organized along the horizontal axis into animal cells and vegetal cells and are further sorted according to their locations along the anterior-posterior axis.

### Model fits reveal a low dimensional geometric description of ascidian gastrulation, consistent with simple effective mechanical dynamics

We first assessed the quality of our model’s fit to three-dimensional light sheet imaging data from a single embryo (ASTEC-Pm1 from Guinard et al.[25]). As described above, spherically curved polyhedral (SCP) tessellations are the only geometries that satisfy our model’s physical and mathematical assumptions. Thus, the extent to which SCP tessellations approximate the cellular geometries observed over time during gastrulation provides a quantitative test of the extent to which ascidian embryos satisfy our model assumptions and thus the validity of the proposed force inference.

Qualitative comparisons at individual time points (Figures 5A-B and 5D-E) suggest that the segmented raw data are remarkably well-approximated by the best-fit SCP tessellations. To assess this quantitatively, we evaluated the local spatial errors in the model’s fit to the data over the entire embryo (Figure 5C and 5F). Globally, the errors are small, on the order of a few percent. The largest systematic error in Figure 5C, on the order of 10%*−*15%, is associated with a single mirror-symmetric pair of posterior mesoderm cells, whose apical surface shapes clearly deviate from uniform curvature. Overlaying the outlines of raw data and model fits (Figure 5F) provides further insights into the nature of the small errors within the bulk of the embryo.

**Figure 5.**
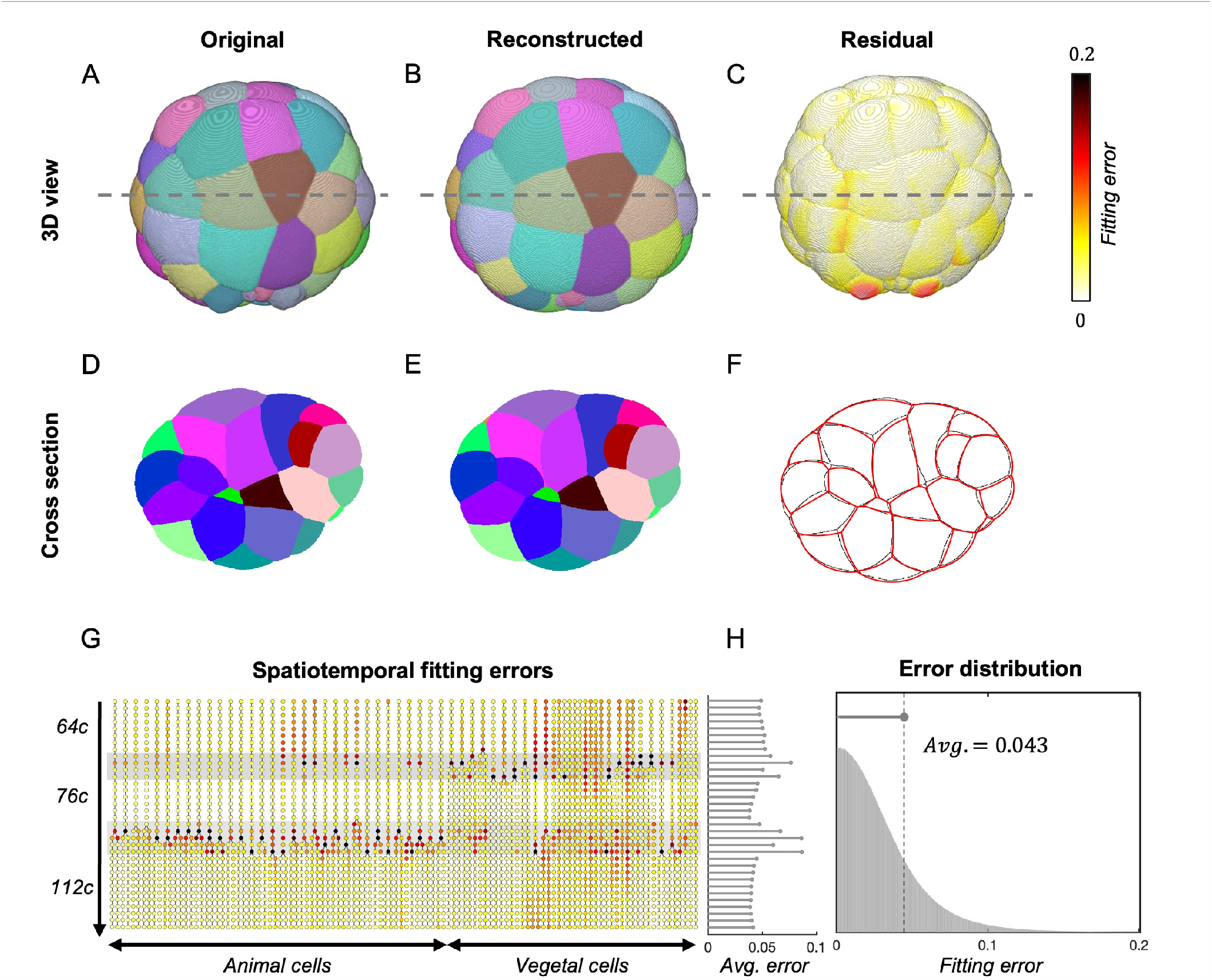
Model fits reveal a low dimensional geometric description of ascidian gastrulation. (A) A three-dimensional view of an ascidian embryo image. Cells are in distinct colors to show the cell boundaries. (B) Reconstructed three-dimensional geometry by best-fitted SCP tessellation. By minimizing the error function, we get the best-fitted geometry parameters Ψ. Then we reconstruct the SCP tessellation using Ψ and visualize it in three-dimensional with the same color scheme as in (A). (C) The residual of membrane pixels deviating from best-fitted SCP tessellation. For each pixel ***r***_*i*_, we compute the deviation |*ϵ*_*i*_ |, normalized by the length-scale of cell size. Then we color the image pixels according to the values of the fitting error and visualize them in three-dimensional. The fitting errors are smaller than 0.1 for most pixels. (D) Two-dimensional cross-section view of the three-dimensional image in (A), visualizing the in-bulk cell shapes. (E) Two-dimensional cross-section view of reconstructed three-dimensional geometry in (B). (F) Comparison of cross-section between original (black) and reconstructed geometry (red). There appears to be a small mismatch. (G) Spatiotemporal fitting errors of ascidian gastrulation shown as lineage map. From top to bottom, each row represents a time point from the 64-cell stage to the 112-cell stage. Each dot represents the cellular-level average of the fitting errors, colored using the same scheme in (C). (H)The distribution of the overall fitting errors (right panel) is close to a half-Gaussian distribution, with the standard deviation equal to 0.043, representing the average fitting error. The left panel presents the average fitting error over time from top to bottom, aligned with the rows in (G). The high average errors appear during the cell-division phase.

These results highlight the precision with which SCP tessellations approximate the ascidian embryo at single time points. To provide a more global assessment, we mapped per-cell approximation errors onto the lineage of the ascidian embryo during early gastrulation, organized by developmental time (from 64 to 112 cell stages) and germ; layer, where the color displayed upon the lineage corresponds to the degree of error, per cell (Figure 5G). Averaging the error across cells in an image, Figure 5H provides a corresponding summary of the distribution of errors at a given time point (indicated by the horizontal dashed line). For most time points the average error is less than 5 percent. Larger errors accumulate during waves of cell divisions as the embryo transitions from having 64 to 76 to 112 cells. The failure of the model to account for the geometries of cells during cell divisions lies in the spherical approximation of its membranes, which fails to capture the dumbbell-like shapes adopted by cells just prior to the computational identification of a new membrane in the segmentation protocol. Resegmenting the original image reduced the fitting errors further, as shown in FigureS4 in SI Part IV.A.

In summary, we find that SCP tessellations provide a highly accurate approximation to the time-evolving geometry of ascidian embryos during early gastrulation. In addition to justifying the use of our proposed three-dimensional force inference scheme, these findings reveal several key features of morphogenesis in early ascidian embryos. First, on timescales relevant for morphogenesis, the geometric dynamics are surprisingly low dimensional. A few geometric parameters, corresponding to simple effective mechanics (see below), suffice to capture the morphogenetic trajectories observed in ascidian embryos. Whereas thousands of pixels delineate the membranes of each cell in raw images, and the three-dimensional computational meshes that represent segmented cell shapes employ hundreds of triangular elements, an SCP tessellation provides a highly accurate representation of an entire embryo with only five parameters per cell.

Second, the close correspondence of observed geometries to SCP tessellations reveals the emergence of effectively simple mechanics from the chemical and mechanical complexities that manifest at shorter length scales and faster timescales. While cells may display complex anisotropic and heterogeneous nonlinear constitutive properties at timescales faster than gastrulation, and length scales smaller than cell-cell contacts [28], at scales relevant to gastrulation, the effective mechanics appear to be surprisingly simple, as embodied in our mechanical model. Scalar values describing isotropic and homogeneous cell pressures, surface and line tensions, which abide by effectively fluid-like balances of normal stresses on individual surfaces and lines of contact, suffice to represent the mechanical state of an entire embryo at each point in time.

Finally, the quality of the data fits suggests that cells within the ascidian embryo always lie close to a static equilibrium of forces since deviations from this would introduce strong velocity-dependent dissipative components to the stresses, resulting in geometries that would be poorly approximated by SCP tessellations. From a physical perspective, the observed dynamics can thus be considered the result of “adiabatic” changes in the configuration of the mechanical state, the morphogenetic timescale being much larger than the mechanical relaxational timescales of the system. From the biological perspective, these results suggest that developmental encoding of morphogenetic dynamics might be ordered in terms of two modes of control: through forms of mechanochemical feedback that act to ensure continuous proximity to static equilibrium, and through developmentally encoded regulatory control over slow changes in pressure, surface tensions and line tensions on individual cells, cell-cell contacts, and junctions.

### A mechanical atlas for gastrulation

#### Choosing values for zero modes

Having established the accuracy and robustness of our force inference scheme for synthetic data, and the quality of our model’s fit to the multicellular geometry of ascidian embryos, we are now poised to construct a mechanical atlas for early gastrulation, Our force inference scheme infers cellular pressures, surface tensions, and line tensions up to the three undetermined global constants (*a, b, c*) defined in equation (2), which cannot be determined from geometry alone. These so-called zero modes reflect the magnitude of ambient pressure (*a*), the absolute scale of force (*b*), and the relative contributions of surface and line tensions (*c*). Crucially, since quasi-static dynamics make the force inference at each timepoint independent of all others, the correspondence between (*a, b, c*) and the physical scales or quantities they represent need not be constant over time. Therefore, we adopt the following approach to visualize and study patterns of relative force across the embryo at each time point and how these patterns change over time.

Without loss of generality, we set the ambient pressure (*a*) to 0 and the force scale (*b*) to 1. This is analogous to the use of arbitrary units (a.u.) in imaging studies where the correspondence between fluorescence and molecular concentration is unknown. To guide our choice of a value for *c*, we note that varying *c* between 0 and its maximum value *c*_*max*_ results in systematic variation in the *relative* values of surface and line tensions on cellular interfaces (Figure 8A). For *c* = 0, surface tensions dominate; for *c* = *c*_*max*_, line tensions dominate; for *c* = *c*_*max*_*/*2, their contributions are approximately equal. However, the *total* contributions of surface and line tensions on each cell - the cellular contractile stress tensor (equation (4)) - remains approximately constant with variation in *c* (Figure 8B). As we discuss further below, this finding reveals a fundamental form of biological flexibility in which it is possible to achieve the same quasistatic force balance through a prescribed infinitude of different combinations of surface and line tensions. Here to aid our visualization of force patterns at individual time points, we set *c* to the intermediate value *c*_*max*_*/*2 where surface and line tension contributions are approximately equal. Finally, to compare forces across time points, we make one additional assumption - that the total hydrostatic energy is conserved over time:∑_*α*_*P*_*α*_*V*_*α*_ ≡ *constant*. We discuss the ambiguity associated with the idea of conservation and the potential biological intuition in detail below.

### A mechanical atlas for gastrulation reveals a rich dynamic correspondence between embryonic axes, the invariant lineage, and patterns of mechanical force

Using these assumptions to fix values for global constants (*a, b, c*), we present a mechanical atlas for ascidian gastrulation, from the 64- to 112-cell stage (Figure 6). Figure 6A shows spatial maps of relative pressures, surface tensions, and line tensions over the entire ascidian embryo at individual (64-cell, 76-cell, and 112-cell) stages, and Figure 6B and 6C map changes in pressure and cellular contractile stress onto the embryonic lineage over the entire interval of early gastrulation. Visual inspection of these maps reveals a striking correspondence between inferred forces and the spatiotemporal patterns of cell identity and cell division encoded in the invariant lineage. Before the first cell division, the deposition of maternal factors, and their relocalization by large-scale cytoplasmic movements [37] establish molecular asymmetries along two embryonic axes: the Anterior/Posterior (AP) axis and the Animal/Vegetal (AV) axis. These asymmetries are elaborated by local cell induction and asymmetric cell division to arrange groups of similarly fated cells along the AP and AV axes. During early gastrulation, these patterns of cell fate are encoded as patterns of force (Figure 6). Surface views of the vegetal hemisphere reveal broad asymmetries in pressure and tension along the AP axis, particularly at the 64-cell and 112-cell stages (first three columns of Figure 6A), while cross-sectional views reveal global force asymmetries across the AV axis (rightmost column of Figure 6A, also quantified in Figure S6C). On a finer scale, we observe strong correlations between patterns of force inferred at individual time points and shared cell identities. This is especially clear in the vegetal half of the embryo where shared patterns of force distinguish all of the major cell types (e.g. endoderm, muscle, notochord, and nerve cord precursors).

**Figure 6.**
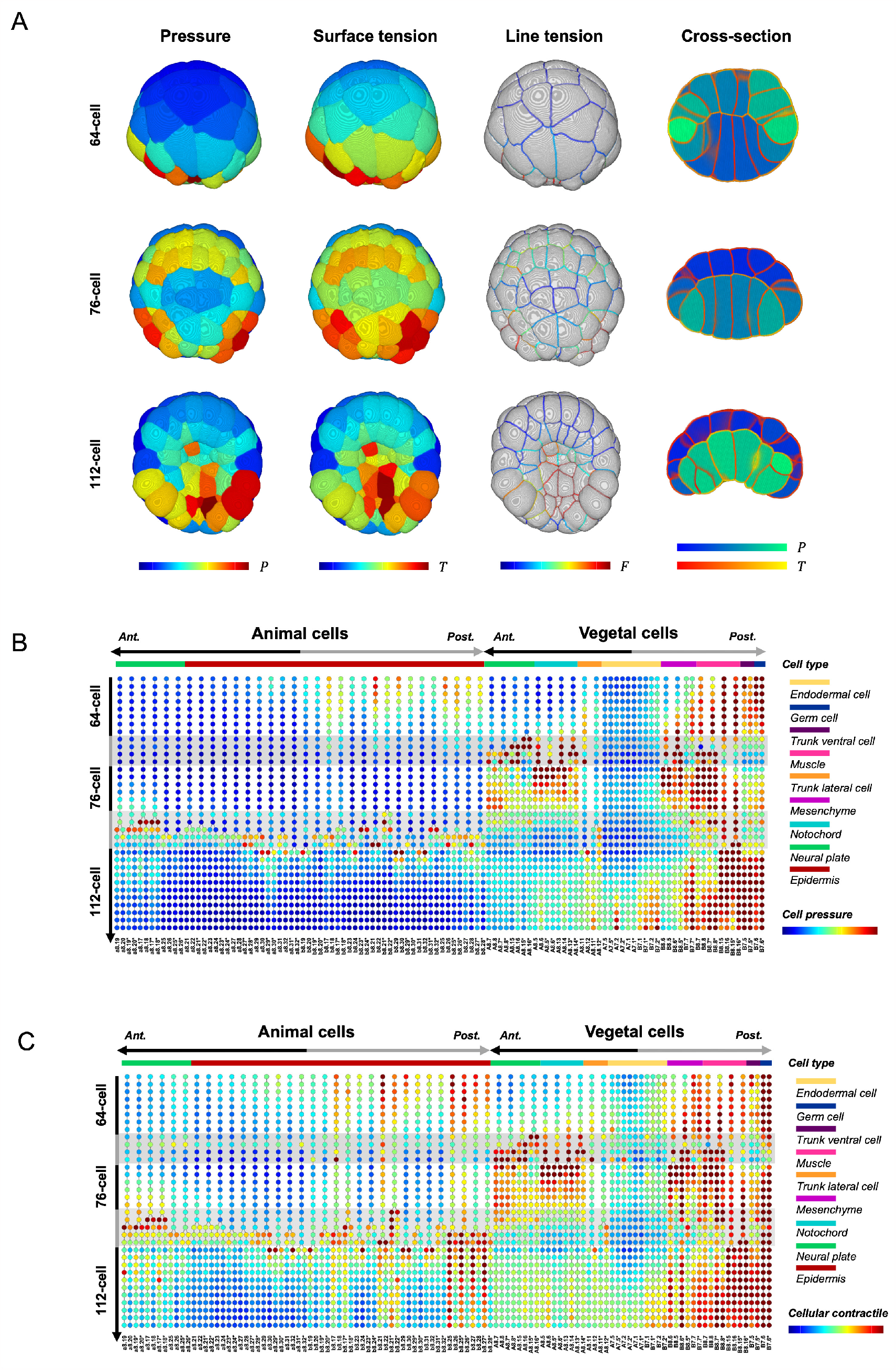
The mechanical atlas for early ascidian gastrulation. (A) Inferred cellular pressures (first column), apical surface tensions (second column), and apical line tensions (third column) in an individual *Phallusia mammillata* embryo (ASTEC-Pm1) at the 64-cell (first row), 76-cell (second row) and 112-cell (third row) stages in vegetal view. In the fourth column, the cross-sectional views present the in-bulk patterns of pressures and surface tensions, using a distinct color scheme from the vegetal visualizations. (B) Lineage map of inferred pressure for gastrulation process in *Phallusia mammillata* embryo (ASTEC-Pm1). Each dot represents a cell at a certain time point, with the color showing the cellular pressure. From top to bottom, each row represents a time point from the 64-cell stage to the 112-cell stage. Cell names are presented at the bottom. Cell types are presented by bars at the top. The cells are organized into animal cells and vegetal cells and are sorted according to locations along the anterior-posterior axis. (C) Lineage map of inferred cellular contractile stress (isotropic part) for gastrulation process in *Phallusia mammillata* embryo (ASTEC-Pm1).

Consistent with the bilaterally symmetric allocation of cell fates, and the bilateral symmetry of cell shapes during early ascidian development [25, 37], the mechanical atlas reveals strong bilateral symmetries in both cellular pressures and membrane/line tensions(Figure 6A; quantified in Figure S6A). We observe slight variations in magnitude between the left and right halves - for example at the 112-cell stage, where the right B7.1-2 and B8.7-8 cells demonstrate higher pressures and apical membrane tensions than the left counterparts. But overall, the regions with relatively high force values are approximately identical. Notably, these overall patterns of force are reproducible across 3 independently imaged embryos (Figure S5C).

Finally, in addition to these spatial patterns observed at individual time points, the mechanical atlas also reveals temporal patterns of force within individual lineages. Consistent with previous work, we observe elevations of pressure and membrane contractile stress just before and during all observed cell divisions. In some lineages (e.g. all animal cells) these forces are transient. In others (e.g. notochord, neural plate, trunk lateral, and mesenchyme), they persist well into the next cell cycle, suggesting that cell divisions may gate transitions in force production. In yet others (e.g. vegetal endoderm, posterior mesoderm, and germ cells) more complex force trajectories unfold between cell divisions.

Altogether, these observations reveal a rich correspondence between dynamically evolving cell states, encoded by patterns of gene expression and signaling, and spatiotemporal patterns of cellular pressure and contractile stress. Given the quality of the model’s fit to the data and its reproducibility across embryos, we conclude that our mechanical atlas captures a robust low-dimensional portrait of mechanical dynamics driving gastrulation a robust signature of the biological control over those dynamics.

### Lineage-specific variation in pressure and contractile stress are highly correlated during gastrulation

Figure 6B and 6C reveal a striking correspondence between spatiotemporal patterns of cellular pressure and membrane contractile stress, which we confirm quantitatively in Figure S7. To examine this further, we considered spatial patterns of surface and line tensions across five distinct types of cell interface - vegetal apical, vegetal lateral, basal, animal lateral, and animal apical (Figure 7A-C). We find that cellular pressures are correlated with heterogeneities in these interfacial surface and line tensions in several ways. Apical membrane tensions are directly correlated with cellular pressures, reflecting the similar curvatures observed across all apical membranes (Figure S6D). Lateral surface tensions are correlated with the mean of the two adjacent cellular pressures (Figure S6E). Surface tensions on basal membranes are correlated with the cellular pressures of the adjacent vegetal cell (Figure S6F). Similar rules appear to govern correlations between pressures and line tensions. For example, line tensions tend to be high on apical junctions between two high-pressure cells (Figure S6G). At lateral and basal junctions, which are formed by three cells, line tensions tend to be high when at least two of the three adjacent cells exhibit high pressures (Figure S6H). For example, at the 112-cell stage, line tensions on junctions formed by two vegetal cells and one animal cell are higher on average than line tensions on junctions formed by one vegetal cell and two animal cells (Figure S6I).

**Figure 7.**
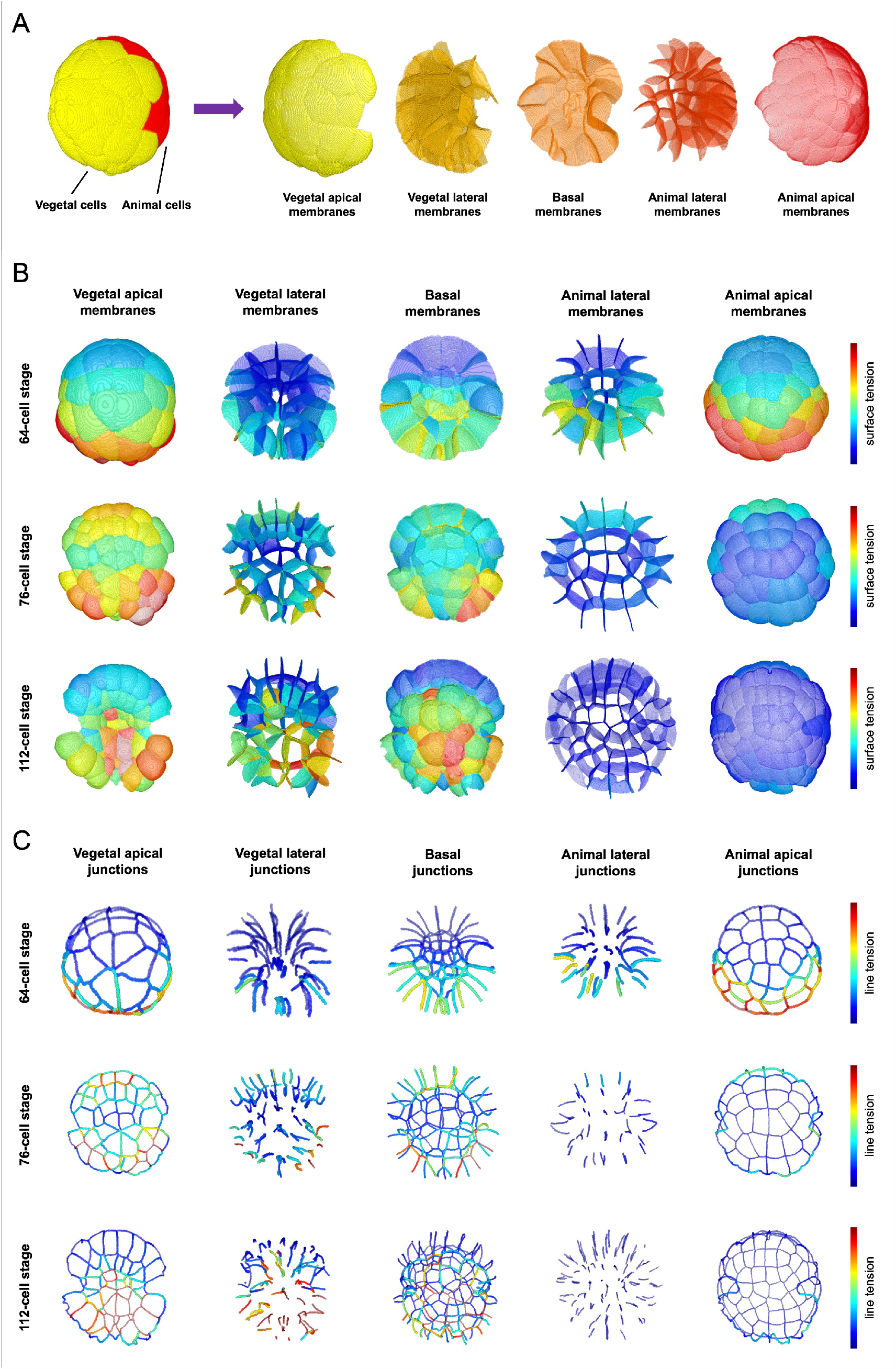
Full mechanical atlas of surface and line tensions for ascidian gastrulation. (A) The ascidian embryo is composed of vegetal cells (yellow, first panel) and animal cells (red, first panel). According to this, the membranes (and junctions) are decomposed into five groups (second panel) - vegetal apical membranes (yellow) that are between vegetal cells and outside, vegetal lateral membranes (dark yellow) that are between vegetal cells, basal membranes (orange) that are between vegetal and animal cells, animal lateral membranes (dark orange) that are between animal cells, animal apical membranes (red) that are between animal cells and outside. (B) Full mechanical atlas of inferred surface tensions in vegetal view, in five layers (five columns) and three stages (three rows). (C)Full mechanical atlas of inferred line tensions in vegetal view, in five layers (five columns) and three stages (three rows).

**Figure 8.**
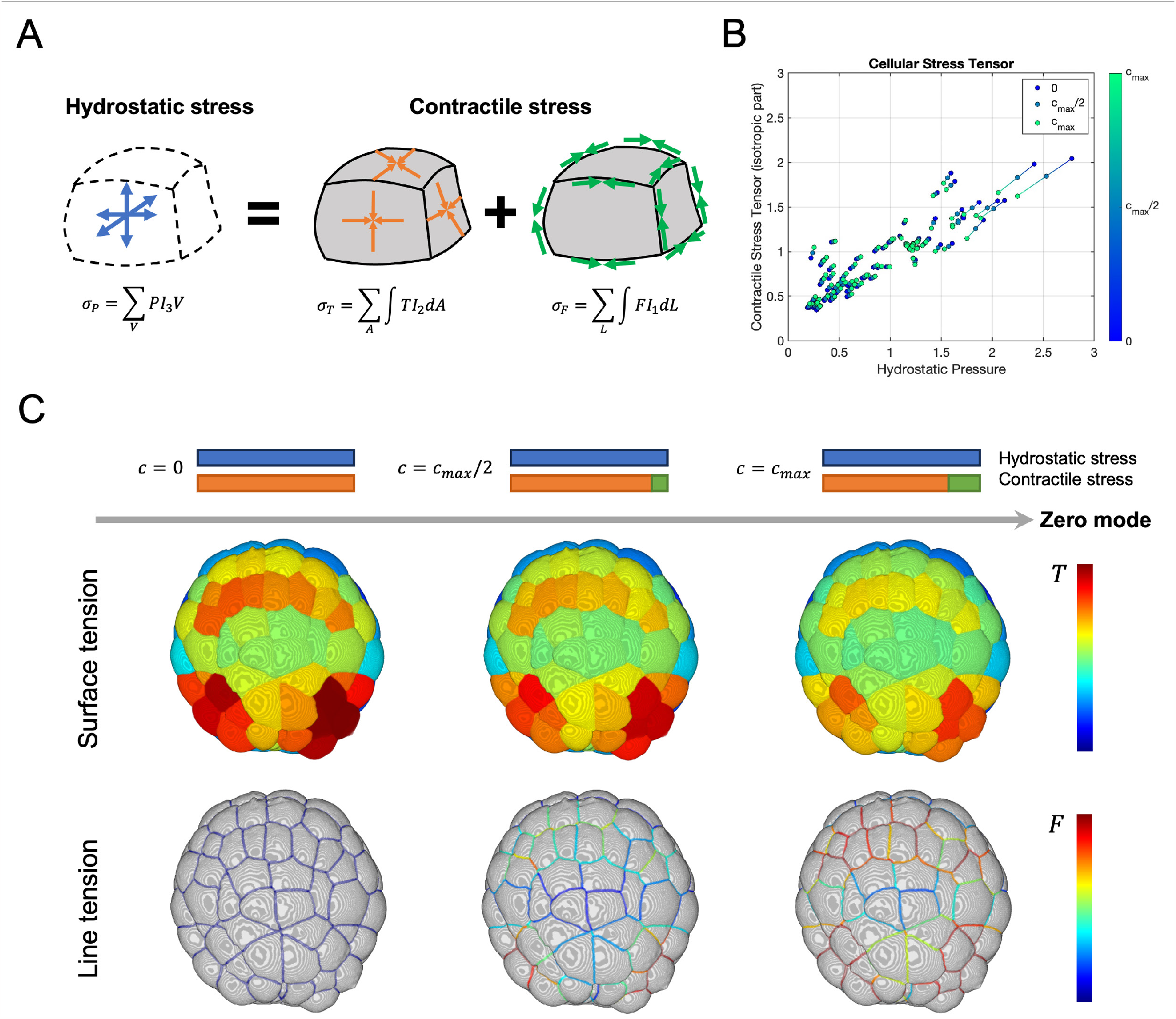
Mechanical flexibility of ascidian embryogenesis. (A) The total hydrostatic stress tensor ***σ***_*P*_ is equivalent to the total contractile stress tensor, which is composed of surface contractile stress ***σ***_*T*_ and line contractile stress ***σ***_*F*_. The zero-mode parameter *c* determines the relative contributions of surface and line contractile stress. (B) The comparison in cellular scale between hydrostatic pressures and isotropic part of contractile stresses – 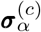 (equation (4)), under different zero-mode parameters *c*. The cellular hydrostatic pressure corresponds with the cellular contractile stress, across all values of *c*. The cellular contractile stresses remain small variance (<20%). (C) The mechanical atlas of apical surface tensions (first row) and apical line tension (second row), under three different values of the mechanical zero-mode *c* (three columns), at a time point in the 76-cell stage. In the left column (*c* = 0), there are zero line tensions and high surface tensions; in the middle column (*c* = *c*_*max*_*/*2), there are moderate line tensions and moderate surface tensions; in the right column (*c* = *c*_*max*_), there are high line tensions and small surface tensions.

The finding that cellular pressures and contractile stresses are correlated is not entirely unexpected. The observation that ascidian embryos operate close to static equilibrium implies that contractile and pressure forces must balance one another globally at all times, while hydrostatic pressure differences, surface tensions, and line tensions must balance locally according to YL laws. Thus the finding that heterogeneities in individual cellular pressures are highly correlated with interfacial surface and line tensions is a non-trivial result. Because cells can independently regulate pressure and local contractile stress, and because these quantities are locally coupled, it remains impossible to infer from geometry alone the mode of regulation. However, by inferring independent spatial and temporal variation in cellular pressure, surface tension, and line tensions at the cellular scale, combining our approach with specific perturbations provides a way to deduce the underlying sources of regulation.

### Global integration of cellular pressure and contractile stress orchestrates ascidian gastrulation

Previous studies of early ascidian gastrulation focussed on how the regulation of myosin II activity in the endoderm cells could drive two distinct phases of endoderm invagination [31]: During the first phase (64-76 cells), apical activation of myosin II accompanies the columnarization and apical constriction of endoderm cells. During the second phase (76-112 cells), circumapical and basolateral activation of myosin II accompanies rapid apico-basal shortening of endoderm cells and invagination of the endoderm plate. 2D vertex models asserting patterns of tension consistent with the observed patterns of myosin II recapitulated the two-phase kinematics of endoderm invagination.

However, while these studies provided valuable insights, they were necessarily limited by a 2D perspective, and by a focus on regulation of myosin II in endoderm cells. Therefore, we used our mechanical atlas to reevaluate the mechanics of endoderm invagination from a global three-dimensional perspective, remaining agnostic to the specific origins of dynamic changes in pressure and tension. Considering the decomposition of interfacial forces in Figure 7, and focussing on endoderm cells, the relative magnitudes of interfacial tensions are broadly consistent with previous work [31, 32]. For example, the ratio of apical::lateral tension in endoderm cells is high at the end of Phase I when myosin II accumulates apically in endoderm cells and they adopt apically constricted columnar shapes, while the ratios of apical::lateral and basal::lateral tension decrease during Phase II as myosin accumulates laterally and endoderm cells undergo apico-basal shortening and invagination.

However, the full mechanical atlas makes it manifestly clear that a much more global integration of lineage-specific changes in pressure and tension across the entire vegetal hemisphere, underlies gastrulation. From the 64-cell to the 76-cell stage, pressure remains low in endoderm cells (A7.1-2, A7.5, B7.1-2). However, pressures increase sharply within anterior notochord and neural plate cells (A8.5-8, and A8.13-16), and remain persistently high within posterior mesoderm cells, such that these cells collectively form a ring of heightened pressure encompassing the lower-pressure endoderm, as the vegetal hemisphere of the embryo flattens at the 76-cell stage. As development progresses to the 112-cell stage, both pressure and surface stress in endoderm cells, a mechanical transformation aligning with the invagination, but they also remain high in the surrounding ring of vegetal cells. Thus, our mechanical atlas reveals a fundamentally new view of ascidian gastrulation involving lineage-specific control over a complex heterogenous pattern of cellular pressure and contractile stress, spanning the entire vegetal hemisphere. These observations highlight the essential importance of developing new approaches to measuring the mechanical stress state of an embryo that go beyond a focus on the forces generated by junctional actomyosin networks.

## Discussion

Efforts to discover design principles for animal morphogenesis have been limited by the challenge of measuring three-dimensional patterns of mechanical force across embryos over time. Here, we have made significant steps towards addressing this challenge. Using a new mathematical theory for three-dimensional cellular aggregates at quasi-static equilibrium, we have identified a low-dimensional predictive map from patterns of subcellular force to equilibrium geometries. Inverting this map yields a robust approach to inferring global patterns of force from segmented light sheet data and identifies the key quantities (zero-modes) that cannot be inferred from geometry alone. Applying this approach to gastrulation in ascidians, we have constructed the first single-cell mechanical atlas of a whole intact embryo executing a major morphogenetic event. The accuracy and robustness with which our model fits observed geometries support a surprisingly simple and low-dimensional view of ascidian morphogenesis in which slow lineage-specific changes in gene expression and signaling govern adiabetic progression through a sequence of quasi-equilibrium states. Contrasting earlier models, our atlas reveals that the global integration of widespread lineage-specific changes in both contractile forces and hydrostatic pressure drives the dynamics of gastrulation. These results highlight the power of our approach to reveal fundamentally new insight into the global dynamics of morphogenesis.

### Comparison to other methods for 3D force inference

Several recent efforts have been made to extend the force inference approach to three dimensions [23, 38, 39, 40]. However, these methods use surface meshes to quantify local geometry; they invoke only local constraints on force balance, and they rely on matrix inverse methods to project high dimensional descriptions of geometry onto a set of possible force arrangements. In particular, current methods fail to encode, mechanically motivated, *global* geometric constraints. The resulting schemes are thus numerically sensitive to noise in the measurement and image analysis protocol, and produce forces that are less accurate and interpretable.

Our approach to 3D force inference differs from these in several fundamentally important ways. First, by making empirically grounded assumptions about mechanical isotropy and homogeneity across cell contacts, and enforcing a global balance of force at equilibrium, we map data to a much lower dimensional space of equilibrium geometries. We infer all possible arrangements of force compatible with the geometries identified by model fits, identify explicitly the remaining degrees of freedom (the zero-modes) that cannot be inferred from geometry alone, and relate these to well-defined physical qualities. Importantly, the quality of data fits provides a robust test of the underlying assumptions and reveals local (in both space and time) failures to meet those assumptions. Our approach is inherently robust in the sense that nearby variants of the same underlying data will map naturally to nearby equilibrium geometries.

### Flexibility in the biological control of embryo geometries

Considering the independent contributions of surface and line tensions [41, 42, 43] allowed us to identify a zero mode that quantifies their relative contributions to producing a given equilibrium geometry. We find that patterns of total cellular contractile stress inferred for a given geometry are insensitive to variation in this zero mode, revealing a fundamental form of biological flexibility in the control of embryo shapes. A growing body of experimental work has shown that embryos can independently regulate contractile force generation on free surfaces, cell-cell contacts, and along specialized (e.g. adherent or tight) junctions [31, 43, 44, 45, 46, 47, 48]. Our results suggest that under different constraints, embryos could use these different modes of regulation to achieve the same morphogenetic outcomes. In the future, it will be interesting to explore whether and how embryos exploit this flexibility to enhance the robustness and/or adaptability of morphogenetic outcomes.

### Ascidian gastrulation follows adiabatic dynamics

The quality of our model’s fit to observed cellular geometries provides strong evidence that ascidian embryos lie always close to a static equilibrium during gastrulation, i.e. the mechanical dynamics are adiabatic. This central finding has nontrivial implications: Given an arbitrary assignment of cellular pressures, surface tensions, and line tensions consistent with our theory, there is no guarantee that there exists an equilibrium geometry for which these forces are balanced. Thus our findings imply the existence of yet-to-be-discovered mechanistic constraints or feedback on force to ensure that ensure continuous proximity to mechanical equilibrium.

### Spatiotemporal variations in pressure and tension drive ascidian gastrulation

An unanticipated feature that the mechanical atlas reveals is that a surprising degree of spatial and temporal variation in cellular pressure underlies gastrulation in ascidians. For several decades, studies of morphogenesis have emphasized a central role in spatiotemporal regulation of actomyosin contractility. Indeed, previous studies identified spatiotemporal patterns of myosin II activation in endoderm cells as a key driver of ascidian gastrulation [31, 32]. While our force inference results do not contradict these earlier findings, they reveal a more complex picture in which stage and lineage-specific spatiotemporal heterogeneities in both pressure and tension drive the morphogenetic dynamics, consistent with a growing list of examples in other systems [26, 27, 40, 49, 50, 51, 52].

Our three-dimensional theory captures physically well-posed and empirically grounded relationships between hydrostatic pressure, volume, surface tension, and line tension that must hold at equilibrium. These relationships make it difficult to infer, from geometry alone, the proximal drivers of heterogeneities in pressure and tension that we observe. Differences in hydrostatic pressure could be determined by changes in osmotic potential within individual cells or e.g. by changes in contractile surface stress driven by changes in myosin II [27, 51, 53]. Similarly, variations in surface and line tensions could be driven by changes in actomyosin contractility or e.g. by the resistance of cell membranes or cell-cell junctions to changes in hydrostatic pressure. An important focus for future work will be to identify the proximal targets for regulation and the associated mechanisms that mediate lineage-specific variations in cellular force.

### Limitations and extensions of the current approach

Despite its success in resolving essential features of ascidian gastrulation, our current approach remains limited in several ways. First, our method fails to identify the scale (in physical units) of force; moreover, the assumption of quasi-static equilibrium precludes the inference of local changes in force magnitude over time, because the scale of force must be determined independently at every timepoint. We note that these limitations apply to any force inference method. Indeed, all force measurements rely on the use of probes to measure a proxy for force (typically displacement or strain), invoking a physical model that relates the proxy measurement to an actual force, and then calibrating the model to produce a force estimate in physical units. Our force inference method uses local cell geometry as the proxy for force and asserts a well-defined physical model for how geometry emerges from the force. As such, it defines a natural approach to calibrating force measurements. In particular, the constraint on global equilibrium implies that a single measurement of force would be sufficient to define global units of force at each time point. Thus in principle, repeated local measurements over time using AFM [12, 13], suction pipettes [10, 11], or embedded oil droplets [16], could provide the required calibration.

Similarly, for the reasons discussed above, our method cannot determine the relative contributions of surface and line tension (quantified by the third zero mode), or how these change over time. However, comparing the change in equilibrium geometry produced by local ablation of individual junctions and individual surfaces and/or whole cells should allow one to estimate the relative contributions of surface and line tension. Although they lie beyond the scope of the present work, these are promising avenues for future efforts.

Finally, we have shown that the assumptions made here regarding mechanical homogeneity, isotropy, and quasistatic dynamics, are well-justified for ascidians. However, it will be important to test whether these assumptions, and our approach, apply to other embryonic contexts, and it will be interesting and important to generalize the approach to account for additional complexities as they arise. For example, it will be interesting to extend the theory of cellular aggregates and the force inference approach to include different types of mechanical anisotropies or inhomogeneities. Similarly, our current theory considers only quasi-static equilibria, ignoring the contributions to force balance coming from effectively viscous or frictional resistances to continuous deformation and flow, which can occur on morphogenetic timescales in some systems [54, 55, 56, 57].

### Outlook for the future

Looking forward, a longer-term scientific goal enabled by the advances laid out in this study is to uncover the regulatory origins of mechanical stresses in embryos and to uncover possible modes of feedback that couple mechanics and regulation. For example, combining a mechanical atlas, as described here, with RNA-based atlases for gene expression at single-cell resolution, and well-defined experimental perturbations, opens the possibility of learning how functional couplings of gene expression and mechanics shape morphogenetic trajectories. Manifestly, the mechanical degrees of freedom are far fewer than the possible regulatory states thus making it possible to study the statistical properties of a highly degenerate genotype-to-phenotype map.

## Data and code availability

The image dataset is available to download and visualize at https://morphonet.org/. The MATLAB codes for the three-dimensional force inference method can be found at https://github.com/siqiliuNU/ForceInferrence.git. The dataset that includes the main results is available at https://data.mendeley.com/preview/8bttsdfstx?a=ec79d745-99c0-41e6-a155-230e31246092.

### Acknowledgements

MM was supported by the National Science Foundation-Simons Center for Quantitative Biology at Northwestern University and the Simons Foundation grant 597491. MM is a Simons Investigator. PL was a senior CNRS researcher. This work was supported by NICHD 1R01HD088831-01 to EMM, a binational “NSF-ANR: Collaborative Research: A mechanical atlas for embryogenesis at single-cell resolution.” (National Science foundation [2204237] and ANR ANR-21-CE13-0046). This project has been made possible in part by grant number DAF2023-329587 from the Chan Zuckerberg Initiative DAF, an advised fund of Silicon Valley Community Foundation.

## Supplementary information

### Part I: Physical model - mechanics determine geometry

#### A. Notations and equations

For a closed-packed cellular aggregate, the cell regions are denoted by *C*_*α*_, *C*_*β*_, …, including the outside region as one ‘cell’. The total number of cells is *N*_*C*_. The contacts between any two adjacent cells are called membrane faces, denoted by *M*_*αβ*_, *M*_*βγ*_, The total number of faces is *N*_*F*_. The junctions between any three adjacent cells are edges of membranes, denoted by *E*_*αβγ*_, …. Each junction is the common edge of three membranes. The total number of edges is *N*_*E*_. In between four adjacent cells, there is a vertex denoted by ***r***_*αβγδ*_. A vertex is a common endpoint of four edges and six faces. The total number of vertices is *N*_*V*_.

Within each cell region *C*_*α*_, there is one isotropic hydrostatic cellular pressure *P*_*α*_. Along each membrane face *M*_*αβ*_, there is one isotropic in-plane surface tension *T*_*αβ*_. Along each edge junction *E*_*αβγ*_, the line tension *F*_*αβγ*_ provides forces in the tangential direction.

Based on the assumptions of mechanical equilibrium, the mechanics determine the geometry of the cellular aggregate by the following force balance equations:

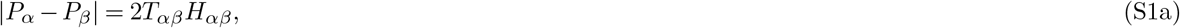

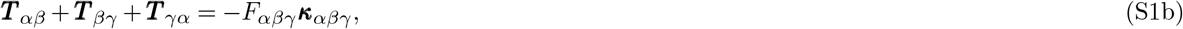

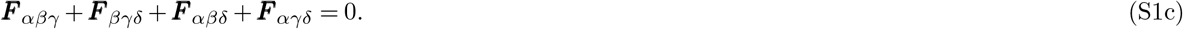

Here *H*_*αβ*_ is the surface mean curvature of *M*_*αβ*_. ***κ***_*αβγ*_ is the curvature vector of the three-dimensional curve *E*_*αβγ*_. The force direction of ***F*** _*αβγ*_ is tangent to *E*_*αβγ*_ at vertex ***r***_*αβγδ*_.

By assumption, the pressures and tensions are homogeneous and isotropic over each cell, each membrane, and each edge. So the membrane shapes are constrained to be spheres (or planes – spheres with no curvature) and the edges are circular arcs (or straight lines – circular arcs with no curvature). Therefore, the equilibrium geometry of a cellular aggregate is polyhedron tessellation or spherically-curved polyhedron (SCP) tessellation. Before parameterizing the two tessellations, we count the dimensionalities of them.

#### B. Dimensionality of a polyhedral tessellation

To count the dimensionality of a polyhedral tessellation, we first investigate the relations between *N*_*C*_, *N*_*F*_, *N*_*E*_, *N*_*V*_ .According to Euler’s characteristic, we have 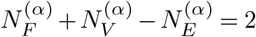 for each polyhedron *C*_*α*_. When we sum the *N*_*C*_ Euler formulas over the tessellation, each face is counted twice, each edge is counted 3 times, and each vertex is counted 4 times. This gives 2*N*_*F*_ + 4*N*_*V*_*−*3*N*_*E*_ = 2*N*_*C*_. Additionally, each edge has two vertices and each vertex has four edges, thus we have 2*N*_*E*_ = 4*N*_*V*_. Using these two relations we derive *N*_*F*_*−N*_*V*_ = *N*_*C*_.

**Claim:** A polyhedral tesselllation of *N*_*C*_ cells has 4*N*_*C*_ degrees of freedom.

**Proof:** The geometry of any polyhedral tessellation can be defined by the positions of all vertices {*r*_*i*_*}*, which have 3*N*_*V*_ degrees of freedom. However, these parameters are not independent owing to the constraint that the faces must be flat. Given the face normal 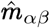, a face edge must be perpendicular to the normal vector, 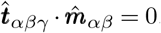. Here 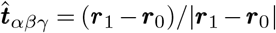 is the edge tangent (Figure S1A). Since each edge corresponds to three faces, there are 3*N*_*E*_ constraints in total. However, these constraints are not independent in two ways. First, for each face, the last edge that closes the face can automatically satisfy the constraint, as shown in Figure S1B. Second, according to the lemma in the next paragraph, there is one additional dependency for each cell. In total, the number of independent constraints is 3*N*_*E*_*−N*_*F*_*−N*_*C*_. The constraints apply to 3*N*_*V*_ degrees of vertices and 2*N*_*F*_ degrees of face normals. Therefore, a polyhedral tessellation has

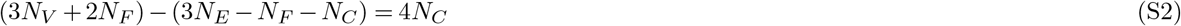

degrees of freedom, using the topology relations 2*N* = 4*N* and *N*_*F*_ *−N*_*V*_ = *N*. □

**Lemma:** For a polyhedral cell *C*_*α*_ in a polyhedral tessellation, the constraints 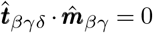 between ‘neighbor’ faces *F*_*βγ*_ and the ‘neighbor’ edges *E*_*βγδ*_ have one dependency. (See Figure S1C.)

**Proof of Lemma:** The constraints of neighbor face normal 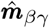 and the three neighbor edges can derive the following relation:

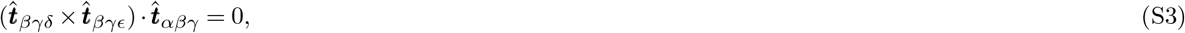

where 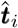 denote the tangent direction of edges. Since 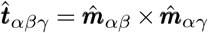. Allowing us to write the relation as

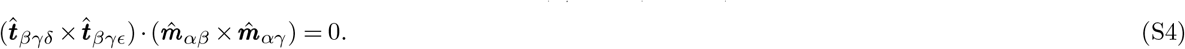

This can be derived as the following equation:

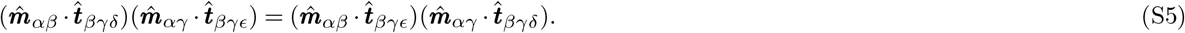

If we multiply all these equations together for neighbor cell pairs *βγ*, there is

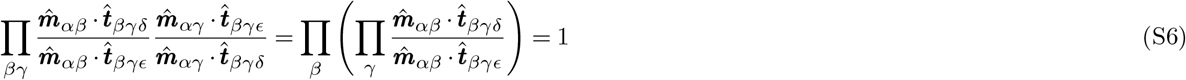

The product in the brackets forms a loop, giving us

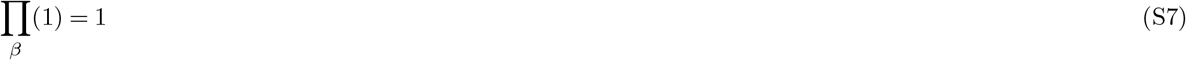

an identity. So these equations have one degree of dependency. Therefore, for each cell, the geometric constraints between neighbor faces and neighbor edges have one dependency. □

#### C. Dimensionality of an SCP tessellation

Similar to polyhedral tessellation, there are geometric constraints in an SCP tessellation because the faces must be spheres and the edges must be circular arcs.

**Claim:** For an SCP tessellation of *N*_*C*_ cells, the dimensionality is 5*N*_*C*_.

**Proof:** The geometry of an SCP tessellation can be defined by vertex locations {***r***_*i*_}, face centroids {***ρ***_*αβ*_}, and face radii {*R*_*αβ*_}. There are 3*N*_*V*_ + 4*N*_*F*_ degrees of freedom in total, but they are under geometry constraints. There are two classes of constraints between vertices and surfaces. First, the distance of a vertex to the face centroid equals the radius (Figure S1D). Since each vertex corresponds to six faces, there are 6*N*_*V*_ such constraints in total. Second, each circular edge is an intersection of three spherical faces, which constrains the three spherical centroids to be colinear along the central axis of the circle (Figure S1E). In total, there are 2*N*_*E*_ such constraints, because each colinearity restricts the other two dimensions. However, these constraints are not independent. According to Lemma 1 in later paragraphs, for each edge, there is one dependency between the two classes of constraints. According to Lemma 2, there is one dependency among the four colinear constraints for each vertex. According to Lemma 3, there is one dependency among all the corresponding colinear constraints for each cell. Therefore, there are *N*_*E*_ + *N*_*V*_ + *N*_*C*_ dependencies among 6*N*_*V*_ + 2*N*_*E*_ total geometric constraints. So the number of independent constraints is 5*N*_*V*_ + *N*_*E*_ *− N*_*C*_. Therefore, an SCP tessellation has

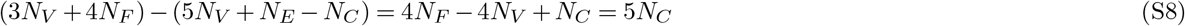

degrees of freedom. □

##### Lemma 1

For an edge *E*_*αβγ*_, the two ends of it are vertices ***r***_1_ and ***r***_2_, as shown in Figure S1F. When |***r***_1_ −***ρ***_*αβ*_ |=|***r***_2_−***ρ***_*αβ*_| and |***r***_1_−***ρ***_*βγ*_ |= |***r***_2_−***ρ***_*βγ* |_are satisfied, |***r***_1_−***ρ***_*γα* |_=| ***r***_2_ −***ρ***_*γα*|_ is satisfied automatically if ***ρ***_*αβ*_, ***ρ***_*βγ*_ and ***ρ***_*γα*_ are co-linear. □

##### Lemma 2

At each vertex ***r***_*αβγδ*_, the centroids of six corresponding faces are coplanar if they satisfy the co-linear constraints, as shown in Figure S1G. Suppose the first three co-linear constraints, corresponding to *χ*_*βγα*_, *χ*_*γδα*_ and *χ*_*δβα*_, are satisfied. These three axes determine a plane, so ***ρ***_*βγ*_, ***ρ***_*γδ*_ and ***ρ***_*δβ*_ can only move in plane. This suggests that there ought to be one constraint instead of two for the last co-linear constraint. The last constraint can be given by

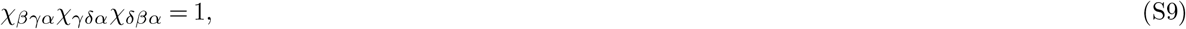

according to Menelaus’ theorem. □

##### Lemma 3

For each cell, if we construct a product of all the Menelaus equations of its vertices,

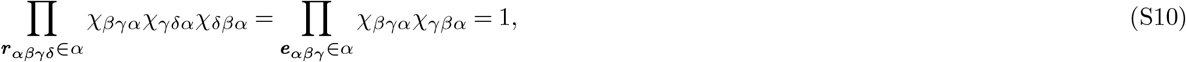

it gives us an identity. That is because *χ*_*βγα*_*χ*_*γβα*_ = 1, and thus all left-hand-side terms cancel each other. □

**Figure S1.**
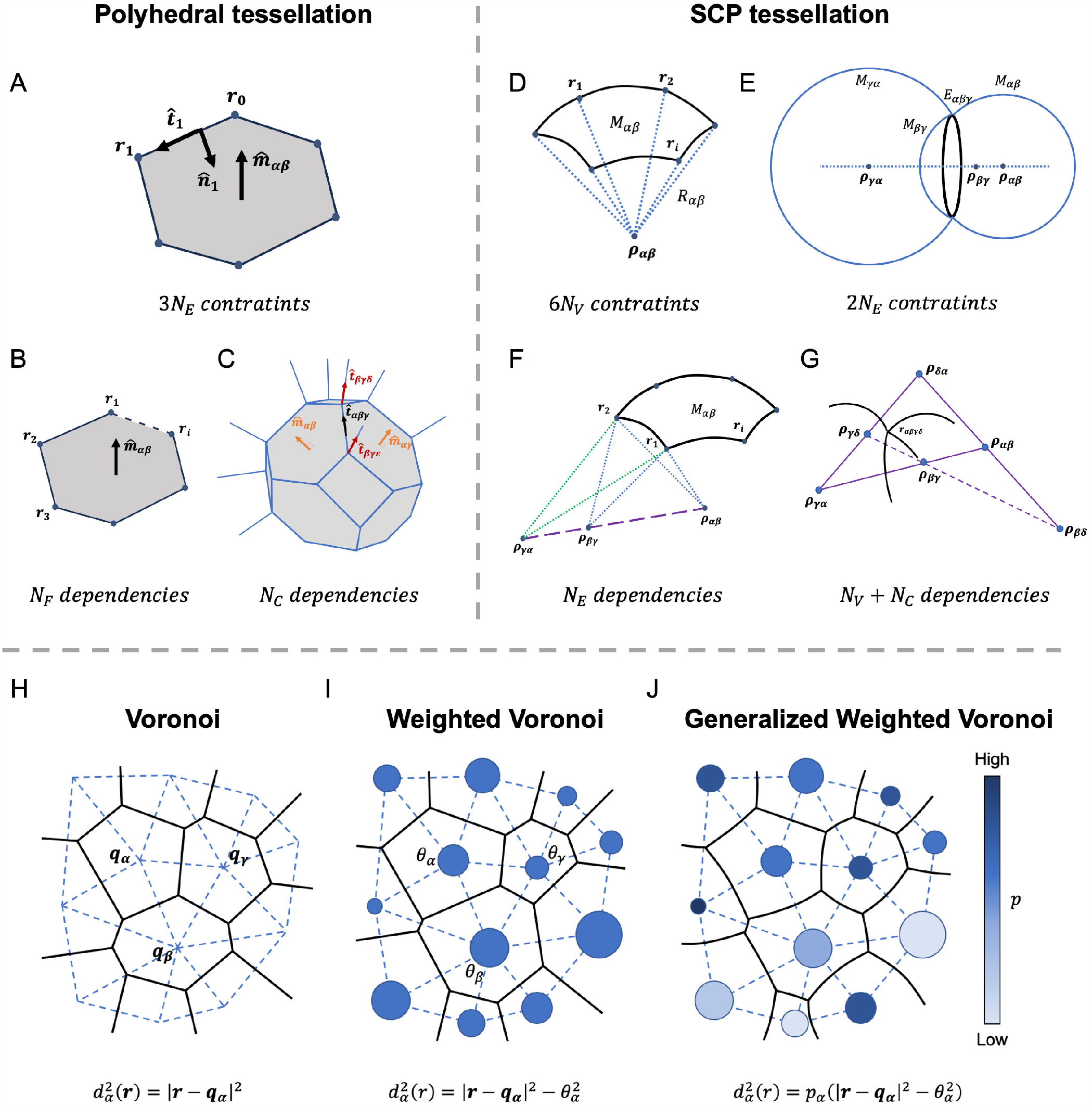
Dimensionalities of polyhedral tessellation and SCP tessellation. (A) Geometric constraints in polyhedral tessellation. (B)(C) Dependencies between geometric constraints in polyhedral tessellation. (D)(E) Geometric constraints in SCP tessellation. (F)(G) Dependencies between geometric constraints in SCP tessellation. (H)(I)(J) The two-dimensional constructions of Voronoi, Weighted Voronoi, and Generalized Weighted Voronoi diagrams.

#### D. Generalized Weighted Voronoi tessellation

To parameterize the 4*N*_*C*_ dimensional polyhedral tessellation and the 5*N*_*C*_ SCP tessellations by independent parameters, we generalize the Voronoi diagram and prove its correspondence to polyhedral and SCP tessellations.

A Voronoi tessellation of *N*_*C*_ cells in three-dimensional space is defined by *N*_*C*_ sites:{*q*_*α*_*α* }. A cell region *C*_*α*_ is a set of points which are closer to ***q***_*α*_ than to other sites: *C*_*α*_ ={***r***|*d*_*α*_(***r***) *< d*_*i*_(***r***), *∀i* ≠ *α* }, and here *d*_*α*_(***r***) =|***r****−****q***_*α*_ | is the Euclidean distance. So the boundary *M*_*αβ*_ between any two neighboring cells is the perpendicular bisector of two corresponding sites, *M*_*αβ*_ ={***r***|*d*_*α*_(***r***) = *d*_*β*_(***r***)}, which is a flat plane. So the shape of a cell is a polyhedron. Therefore, Voronoi tessellation is a subspace of the Polyhedral tessellation space, whose dimensionality is 3*N*_*C*_. Figure S1H shows the two-dimensional case of Voronoi tessellation.

Now, we can modify the definition of distance by subtracting (or adding) a parameter 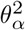 to the squared Euclidean distance,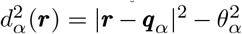. A cell region *C*_*α*_ is defined in the same way: *C*_*α*_ ={***r***|*d*_*α*_(***r***) *< d*_*i*_(***r***), *∀i* ≠ *α* }. We call this class of tessellation a Weighted Voronoi tessellation, where *θ*_*α*_ is the weight for each cell *α*. As we proved below, cell shapes in Weighted Voronoi tessellation are polyhedrons. Since the dimensionality of Weighted Voronoi tessellation space is 4*N*_*C*_, it is equivalent to the space of Polyhedral tessellation. Therefore, we can use the independent parameters 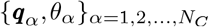 to parameterize Polyhedral tessellation. Figure S2I shows the two-dimensional case of Weighted Voronoi tessellation.

**Claim:** Cell shapes in Weighted Voronoi tessellation are polyhedrons.

**Proof:** For any two points *r*_1_ and *r*_2_ at the two-cell boundary *M*_*αβ*_ satisfy

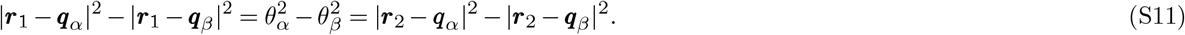

Algebraic simplification yields

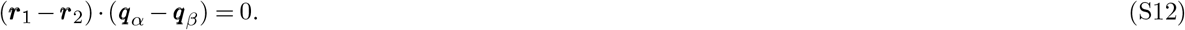

Thus ***r***_1_*−****r***_2_ is perpendicular to ***q***_*α*_*−* ***q***_*β*_. Since the two points are randomly chosen on the boundary, we can derive that the boundary face is an orthogonal plane to the line *q*_*αβ*_. Therefore, the shapes of cells are polyhedral. □

Now, we can further modify the definition of distance by a multiplicative factor *p*_*α*_ to obtain 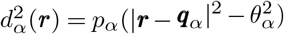 and construct the tessellation in the same way. We call such tessellation a Generalized Weighted Voronoi (GWV) tessellation, and we call *p*_*α*_ as the power of cell *α*. The dimensionality of GWV tessellation space is 5*N*_*C*_, determined by independent parameters 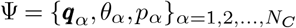. As proved below, the shapes of cell boundaries are spherical, thus a GWV tessellation is an SCP tessellation. Since the dimensionalities of GWV and SCP are equal, there exists a one-on-one map between the two spaces. Therefore, SCP tessellation can be parameterized by Ψ. Figure S2J shows the two-dimensional case of GWV tessellation.

**Claim:** The shape of any two-cell boundary is spherical in a GWV tessellation.

**Proof:** Any point *r* at boundary *M*_*αβ*_ satisfies 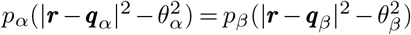. Multiplying this equation by (*p*_*α*_ *− p*_*β*_) and simplifying yields

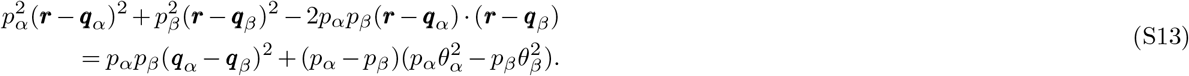

The left-hand side of the above equations is a square, while the right-hand side is a constant independent of ***r***. So this equation can be further simplified as

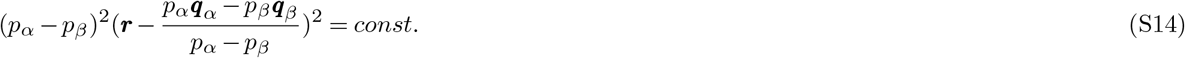

Defining

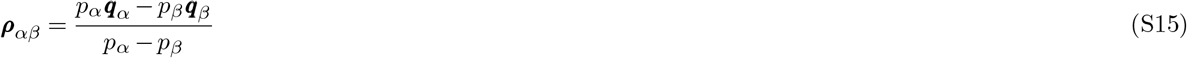

and

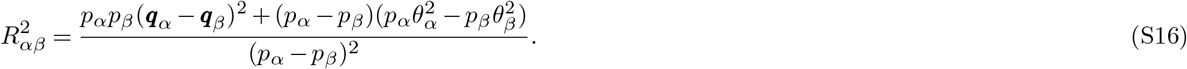

the equation for ***r*** becomes |***r****−****ρ***_*αβ*_ |^2^ = *R*_*αβ*_, which describes a sphere with centroid ***ρ***_*αβ*_ and radius *R*_*αβ*_. So the shape of the boundary *M*_*αβ*_ is a spherical section. □

In summary, consider the intrinsic geometric constraints in polyhedron tessellation or SCP tessellation, the two types of tessellation have 4*N*_*C*_ and 5*N*_*C*_ degrees of freedom, respectively. Polyhedron tessellations can be constructed by independent parameters of Weighted Voronoi tessellation, and SCP tessellations can be constructed by independent parameters of GWV tessellation.

### Part II: Inverse method - infer mechanics from geometry

#### A. Integral of vector area by divergence theorem

Suppose the geometry parameters of SCP tessellation are given by 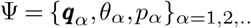, we are able to solve the inverse problem that infer mechanical values *P*_*α*_, *T*_*αβ*_, *F*_*αβγ*_ from Ψ. Before investigating the analytical inverse mappings, we first introduce the concept of vector area and its integral on a closed shape.

For a flat surface in three-dimensional space, the vector area is a vector 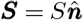 with the face area *S* as the magnitude and the face normal 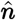 as the direction. For a face element *dS* on a curved face, the vector area is 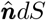*dS*. Consider a closed three-dimensional shape (Figure S2A), the total vector area over the surface is zero according to the divergence theorem, as proved below.

**Theory:** For any closed three-dimensional shape *V*, the integral of the vector area 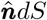*dS* over the closed shape surface *∂V* is zero:

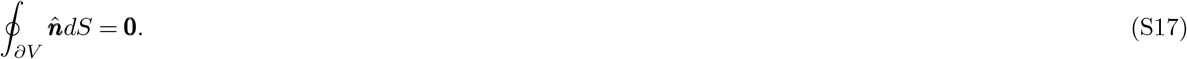

**Proof:** According to the three-dimensional divergence theorem, for any vector of matrix field *F*, there is 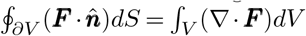. Taking ***F*** as the identical matrix ***I***, we will get 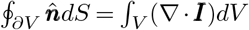. Since Δ·***I*** = **0**, the right-hand-side of the equation is zero. □

**Corollary:** For any closed two-dimensional shape *A*, the integral of the edge normal vector 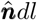*dl* over the closed shape boundary *∂A* is zero:

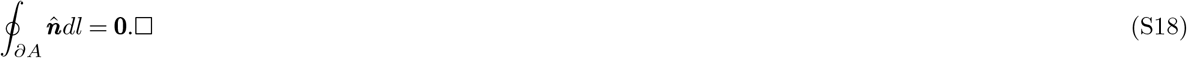

#### B. Two-dimensional mechanical dual graph

We first introduce the two-dimensional analog of the inverse problem. In a two-dimensional cellular lattice, the equilibrium geometry under uniform pressure is a polygonal tessellation that can be parameterized by a two-dimensional Weighted Voronoi diagram. For any membrane *M*_*αβ*_, the line connecting two corresponding Voronoi sites ***q***_*α*_ and ***q***_*β*_ is perpendicular to the membrane. Therefore, we call the triangular lattice formed by Voronoi sites a ‘dual graph’ to the polygon tessellation. As shown in Figure S2D and S2G, consider the dual triangle of the vertex ***r***_*αβγ*_ and apply the divergence theorem, we have

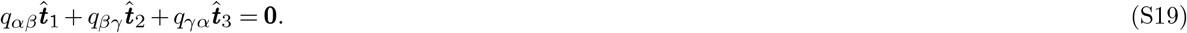

Here, 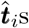 are the tangent direction of edges, and equivalently they are normal directions of dual lines. This equation have the same form and coefficients - the 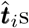 - as the tension balance equation:

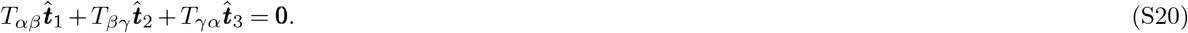

Therefore, the value of membrane tension is proportional to the dual lattice length. Thus we get the analytical solution of tensions:

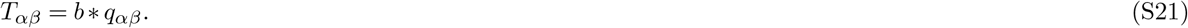

Here the undefined constant *b* represents the scale of tensions, and it is a global constant across the entire tessellation. In this way, we construct the correspondence between the forces and the vector areas of the dual graph, allowing us to get the analytical mapping from geometry parameters to force values.

Now consider the equilibrium geometry of Circular Arced Polygon (CAP) tessellation under non-uniform pressures. This can be parameterized by a two-dimensional GWV tessellation, taking 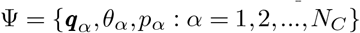 as parameters. Now the membranes are curved, thus we need a new construction of the dual graph at each vertex and along each membrane. For any point ***r*** on the membrane of cell *C*_*α*_, we define the dual point with respect to ***r*** as

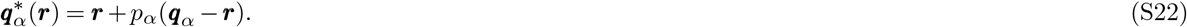

**Figure S2.**
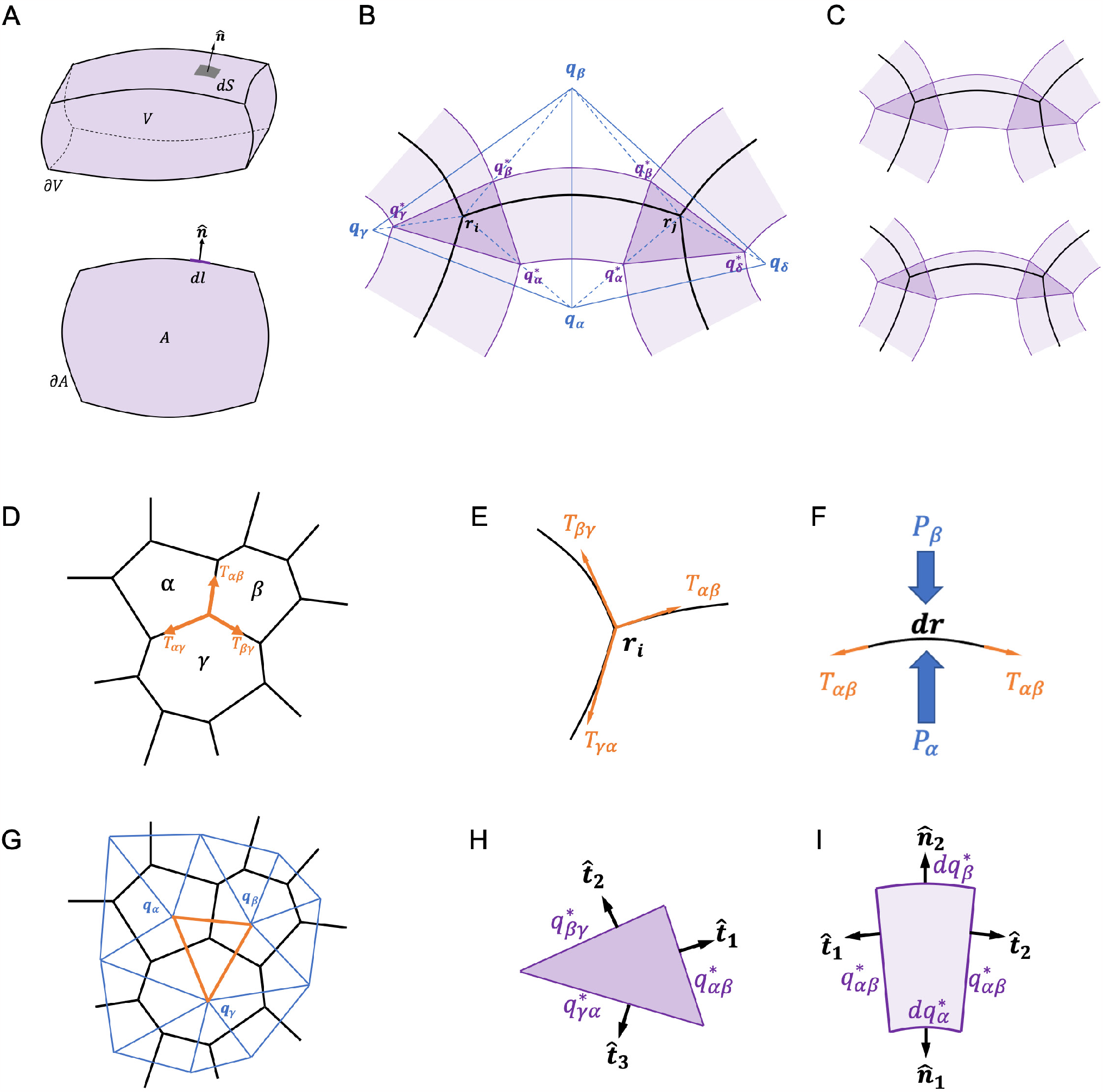
Integral of vector area and two-dimensional mechanical dual graph. (A) Integral of vector area in three-dimensional and two-dimensional cases. (B) The construction of a two-dimensional dual graph. (C) Different choices of dual graph construction in the same CAP geometry. (D)(G) The force balance relation and the corresponding vector areas in a dual graph of polygonal tessellation. (E)(F)(H)(I) The force balance relations and the corresponding vector areas in a dual graph of CAP tessellation.

As shown in Figure S2B, corresponding to the membrane *M*_*αβ*_, the two dual points form the dual line 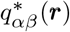. At a vertex, the three dual lines form a dual triangle; along the membrane, the dual line swipes a curved quadrilateral shape.

**Claim:** The dual line 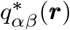 with respect to the point ***r*** at membrane *M*_*αβ*_ is perpendicular to the membrane at ***r***, and the length is a constant along the membrane.

**Proof:** The center of the membrane circular arc is ***ρ***_*αβ*_ = (*p*_*α*_***q***_*α*_ *− p*_*β*_ ***q***_*β*_)*/*(*p*_*α*_ *− p*_*β*_). Consider the vector 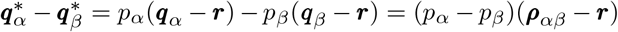, it is parallel to ***r*** *−* ***ρ***_*α*_. This parallel is equivalent to the perpendicularity between the dual line and membrane. On the other hand, from the equation we have 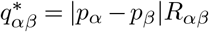. So the length of the dual line is a constant along the membrane. □

Therefore, we can apply the divergence theorem to the dual triangle at the ***r***_*i*_ (Figure S2E and S2H), which gives us:

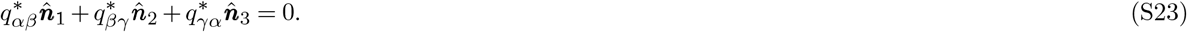

Because of the orthogonality between dual lines and membrane, this equation corresponds to the membrane tension balance equation. So the tension is proportional to the length of the dual line:

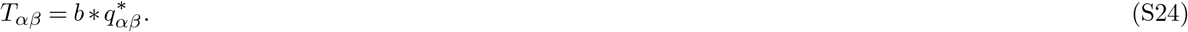

Similarly for the dual curved quadrilateral corresponding to a membrane element *dr* (Figure S2F and S2I), applying the divergence theorem derives

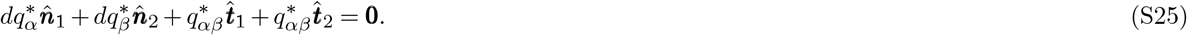

Here, 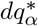 and 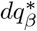 are the length which is swiped by ***r*** along 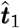 and 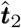 are tangent directions of the edge; 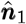 and 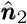 are normal directions of the edge. This has the same form and coefficients as the elemental pressure tension balance equation:

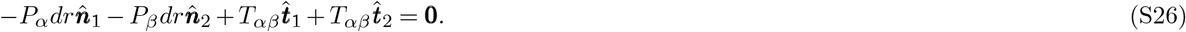

Therefore, the solution of pressure is

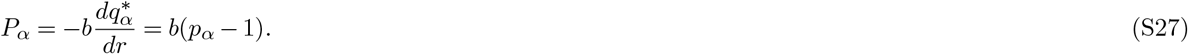

However, we notice that the dual graph is not a unique choice. If we set the rescaled dual points of ***r*** as 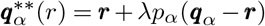, by the free parameter *λ*, then the dual graph is different (Figure S2C). Compared to the previous dual graph, the edge lengths of the quadrilateral have changed. Thus the analytical solution of forces is

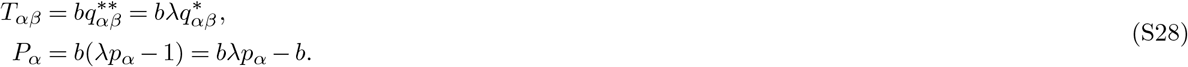

Reset the constants (*b, λ*), we have the general form of analytical solution of mechanics:

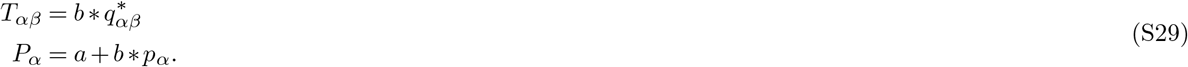

The undefined constants (*a, b*) can be set as any value and will not affect the equilibrium. The physical meaning of *a* is the background pressure and *b* is the mechanical scale.

#### C. Three-dimensional mechanical dual graph

In the three-dimensional case, we can construct the dual graph in a similar way. For the polyhedral tessellation, the dual lattice is formed by directly connecting the Voronoi sites. For any membrane face *M*_*αβ*_, the corresponding dual line *q*_*αβ*_ is perpendicular to the face. Thus at an edge *E*_*αβγ*_, the three dual lines form a dual triangle, which is orthogonal to the edge. Equivalently, the tangent direction of the junctional edge 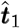 is the normal direction of the dual triangle. As shown in Figure S3A and S3E, applying the two-dimensional divergence theorem to this triangle, we get the equation 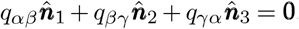. Comparing to the surface tension balance equation 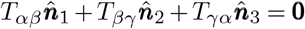., the values of membrane surface tensions are proportional to the length of dual lines:

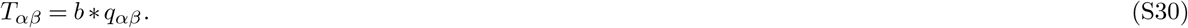

Here *b* is an undefined global constant, representing the scale of surface tensions. For a vertex ***r***_*αβγδ*_, the four dual triangles form a dual tetrahedron. Applying the three-dimensional divergence theorem to the dual tetrahedron, we get

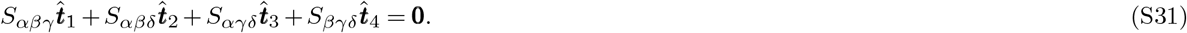

Here the 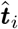and *S*_*i*_ are normal directions and areas of the dual triangle. This equation has the same form and coefficients - the 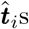 - as line tension balance equation (S1c). Therefore, the values of line tensions are proportional to the corresponding dual triangle areas:

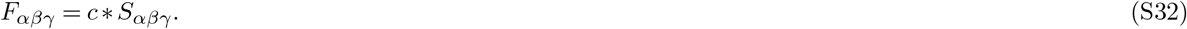

Here *c* is another global constant that represents the scale of line tensions. In summary, (S30) and (S32) provide the analytical solutions of membrane and surface tensions given by Voronoi parameters. We notice that line tension balance and surface tension balance are decoupled with each other, thus the two scales (*b, c*) are decoupled in the solution.

Now, let us consider the SCP tessellation and the construction of the dual graph. As in the two-dimensional scenario, we define the dual point 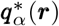 with respect to the point ***r*** on membrane *M*_*αβ*_ by equation (S22). By the same proof, the dual line 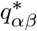 is perpendicular to the membrane face at ***r*** and the length is constant along the membrane. Additionally, for an edge *E*_*αβγ*_, the dual triangle has a constant area

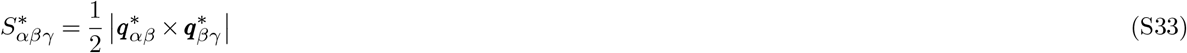

along the edge. Also, the dual triangle is orthogonal to the edge at *r*. Therefore, at a vertex, the four dual triangles form a dual tetrahedron; along an edge, the dual triangle swipes a curved triangular tube; over a membrane face, the dual line swipes a shell with a certain thickness. These three dual graphs correspond to the three force balance equations (S1a) *−*(S1c), as shown in Figure S3B-D and S3F-H.

First, consider the dual tetrahedron at the vertex ***r***_*αβγδ*_, the four vector areas satisfy

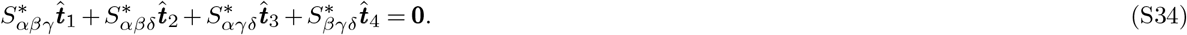

This equation corresponds to the line tension balance equation (S1c). So the solution of line tensions in this case is:

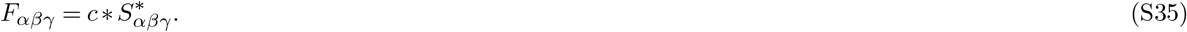

Here *c* is a global constant across the entire tessellation.

Second, consider the elemental dual triangular tube with respect to *dr* along the edge *E*_*αβγ*_, the five vector areas satisfy

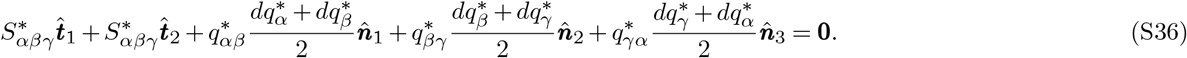

Here 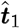 and 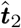 are tangent directions of the edge at two ends. The 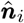 are normal vectors of lateral faces. Note that the lateral faces are trapezoids because the tube is curved. This equation corresponds to the elemental version of force balance equation (S1b), expressed as

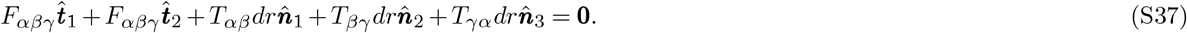

Therefore, applying the line tension solutions and comparing the terms of two equations, the solution of the surface tension is

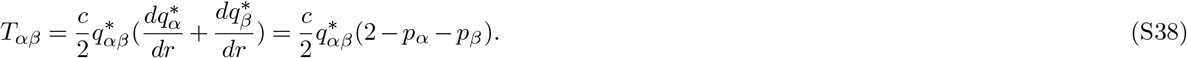

Third, consider the dual squared shell with respect to a small surface element *dr*_1_*×dr*_1_ on the membrane *M*_*αβ*_, the six vector areas satisfy

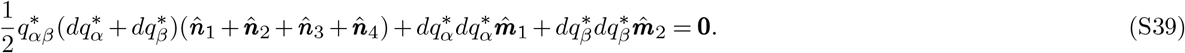

Here 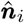 are lateral face normals and 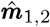 are membrane face normal vectors. This equation corresponds to the elemental version of YL equation (S1a), expressed as

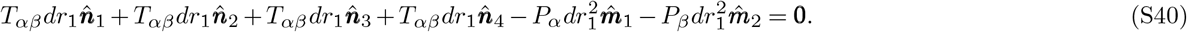

Therefore, applying the surface tension solution and comparing the equation terms, the solution of cellular pressure is

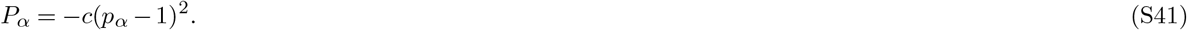

Using a similar trick as in two-dimension to rescale the dual graph, we can derive the general form of analytical solutions of mechanics:

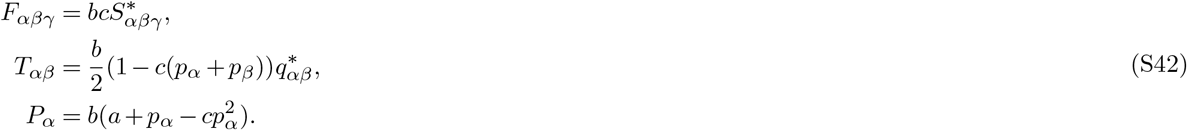

In this solution, the global constants (*a, b, c*) are undefined free parameters.

#### D. Mechanical zero-modes

The undefined constants (*a, b, c*) imply that the inverse mapping from geometry to mechanics is not unique. Different mechanical states can determine the same geometry of a cellular aggregate. The mechanical states allow three classes of variation that can result in the same geometry, which are three mechanical zero-modes. The first class of variation with respect to *a* represents the ambient pressure scale. The second class of variation with respect to *b* represents the scale factor of all forces. The third class of variation with respect to *c* represents the relative contribution of contractility of line tensions.

Let us consider a simple two-cell example to facilitate a physical understanding of the three mechanical zero-modes (Figure S3I and S3J). To set the scene, the two-cell system is defined by the following geometry parameters as: ***q***_1_ = [1, 0, 0], ***q***_2_ = [*−*1, 0, 0], *p*_1_ = *p*_2_ = 1, *θ*_1_ = *θ*_2_ = 2. And also the background parameters: ***q***_0_ = [0, 0, 0], *p*_0_ = 0, *θ*_0_ = 0. This results in two spherical membranes and a flat membrane in the middle. The centroids and radii of three membrane faces are: ***ρ***_10_ = ***q***_1_, ***ρ***_20_ = ***q***_2_, *R*_10_ = *R*_20_ = 2; ***ρ***_12_ = *∞, R*_12_ = *∞*. In order to do the inverse mapping to mechanics, we take a point 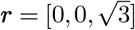 on the edge to construct the dual graph. The dual points are: 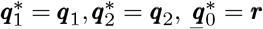. Then the lengths of the dual lines and the area of the dual triangle are: 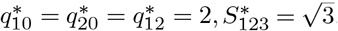. So the forces are:

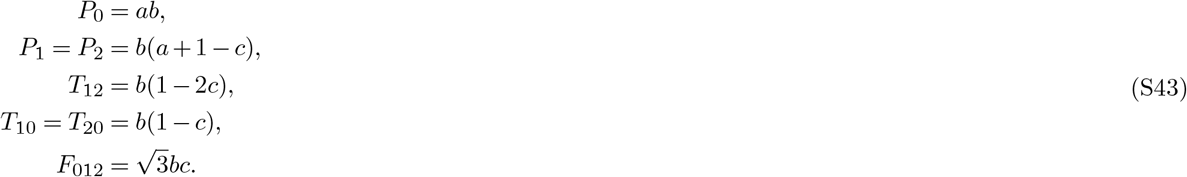

Now, let us investigate the three mechanical zero-modes in detail. First, if we vary the constant *a*, all pressures will change by the same value *ab*, but the pressure differences remain unchanged. Second, if we vary the constant *b*, all the forces will change their physical units, but the relative values of the forces remain unchanged. Third, as for the constant *c*, it changes the relative value of surface tensions. If *c* = 0 (non-line-tension case), the three membrane tension are equal *T*_10_ = *T*_20_ = *T*_12_ = *b*. If *c* increases (e.g. *c* = 0.1), then the middle surface tension will decrease relative to the other two surface tensions *T*_10_ = *T*_20_ *> T*_12_. At the same time, the relative value of pressures will change as well.

By this example, we also notice that the parameter *c* can not be two large. In our assumptions, both surface tensions and line tensions are contractile forces, indicating that the solutions of *F* and *T* must be positive. This confines the permissible choice of parameter *c* that there should be *c*≥0 and *c*≤*c*_*max*_ = max_*αβ*_ (*p*_*α*_ + *p*_*β*_)^*−*1^.

The three constants (*a, b, c*) are global parameters when the cellular aggregate is close-packed. If the cells belong to more than one connected set, then the constant *b* could be different for different sets. Another exception that the parameter *c* is not a global constant is the following example. Consider the linearly connected cells with the same pressure (Figure S3K). For each circular edge between two adjacent cells, the line tension can be any values that are within the permissible range. Each surface tension of the membranes between two adjacent cells can be determined according to the line tension of its edge respectively. That is because the cells are not close-packed, thus there are fewer constraints between line tensions and surface tensions.

**Figure S3.**
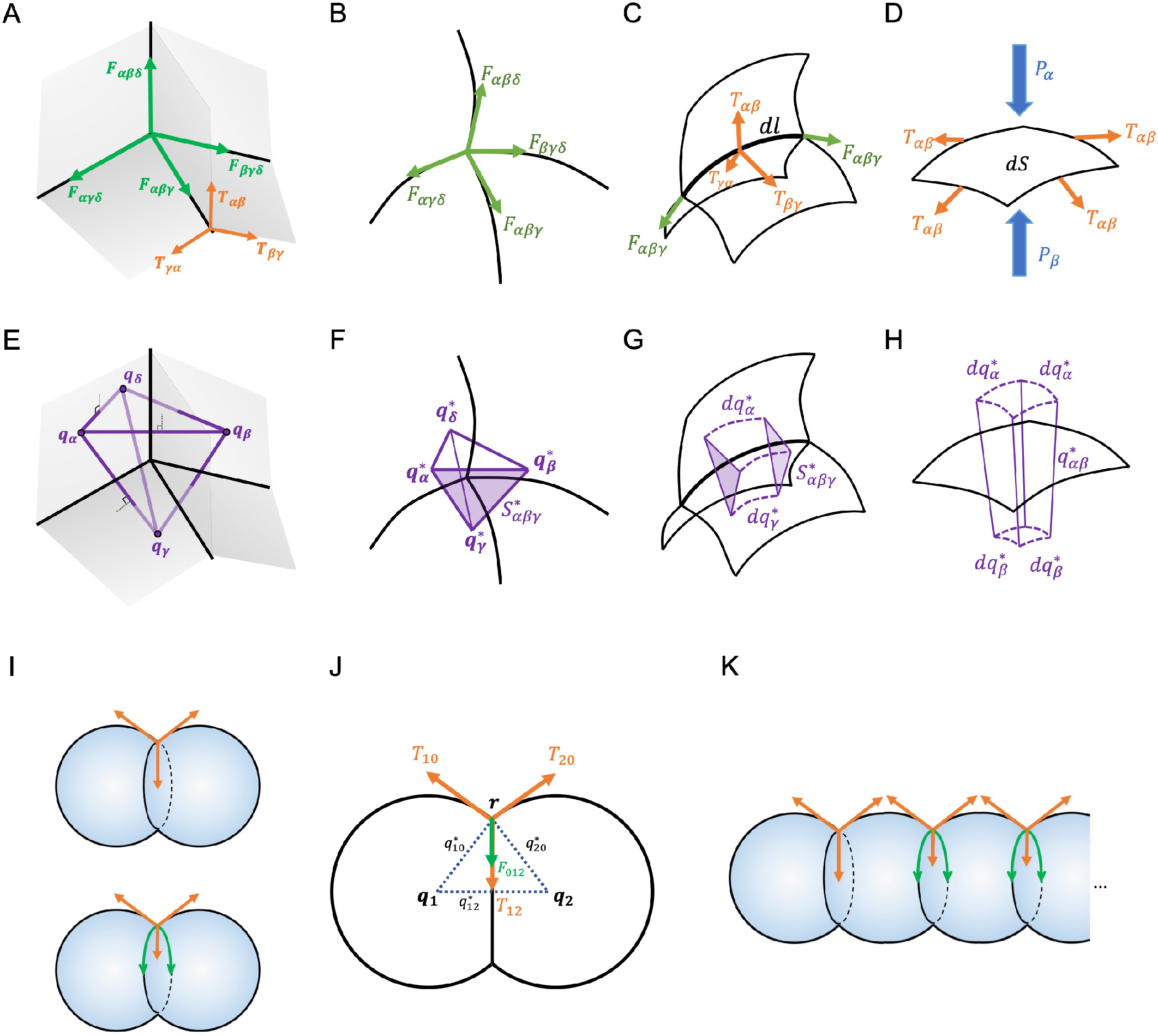
Three-dimensional mechanical dual graph and mechanical zero-modes. (A)(E) The force balance relations and the corresponding vector areas in a dual graph of polyhedral tessellation. (B)(C)(D)(F)(G)(H) The force balance relations and the corresponding vector areas in a dual graph of SCP tessellation. (I) A two-cell example representing the third mechanical zero-mode. (J) The cross-sectional view of (I) and notations. (K) An example of non-close-packed cellular aggregates with non-global choices of the third mechanical zero-mode.

#### E. Stress tensor

In order to coarse-grain the mechanics at the cellular level, we introduce the definition of the cellular stress tensor. Inside a cell, the cell volume is under an isotropic stress, pressure, ***σ***_*P*_ = *P*_*α*_***I***_3_ (where ***I***_3_ denotes the identity matrix in three-dimensions). On a membrane, surface tension generates an in-plane mechanical stress ***σ***_*T*_ = *T*_*αβ*_ ***I***_2_ (where ***I***_2_ is the in-plane projection of identity matrix). Along each edge, line tensions generates a tangential stress ***σ***_*F*_ = *F*_*αβγ*_ ***I***_1_ (where ***I***_1_ is one-dimensional projection of identity matrix). Accounting for all of these, the cellular stress tensor is defined as

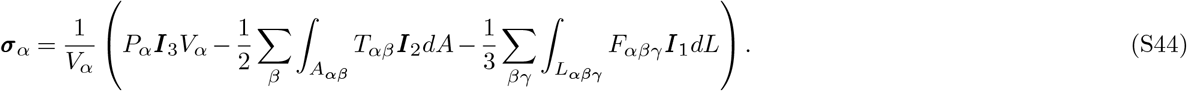

Here the 1*/*2 and 1*/*3 factors represent the share of the stress on boundaries by two or three cells. The minus signs before the second and third terms denote that the surface stress and line stress both contribute contractility of *σ*_*α*_, which can be separately expressed as the cellular contractile stress tensor:

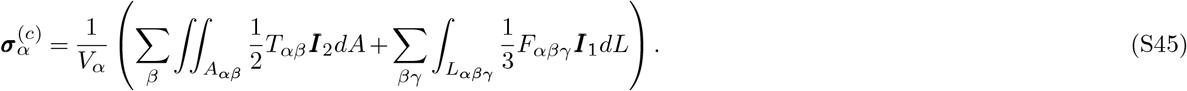

By construction, the cellular stress tensor is a 3*×*3 symmetric tensor, because ***I***_1_, ***I***_2_, ***I***_3_ are all symmetric tensors. The dimension of the stress tensor is [*σ*] = *F/L*^2^, while the dimensions of three mechanical inputs are [*P*_*α*_] = *F/L*^2^, [*T*_*αβ*_] = *F/L*, and [*F*_*αβγ*_] = *F*.

The cellular stress tensor has both isotropic and anisotropic components. The isotropic part can be straightforwardly quantified by the trace of the matrix, 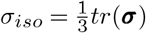. The traceless anisotropic part can be quantified by the scalar shear stress termed von Mises Stress, 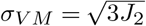, through the second characteristic *J*_2_ of the deviatoric matrix:

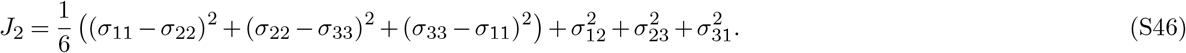

### Part III: Numerical Scheme - fit geometry to image

#### A. Least squares fitting on membrane pixels

In order to quantify the SCP geometry from a three-dimensional image of a cellular aggregate, we are going to find the GWV parameters 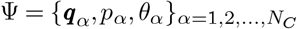 by least squares fitting. Given an image that is segmented according to the cell shape, we get a set of coordinates of membrane pixels *r*_*i*_ and the two cell indices *α, β* that each pixel belongs to. Then the deviation of each pixel to the theoretical sphere is

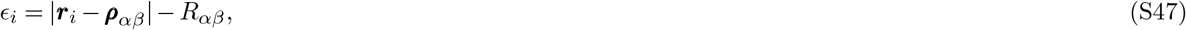

where the sphere centroid ***ρ***_*αβ*_ and radius *R*_*αβ*_ are functions of Ψ, determined by equation (S15) and equation (S16). Therefore, the global mean-squared-deviation (MSD) function is given by

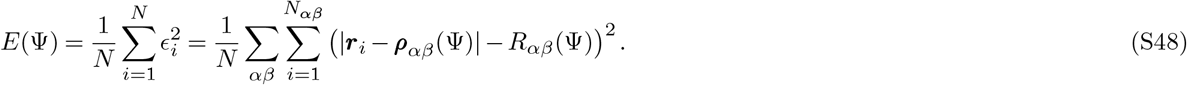

By minimizing the MSD function, we could get the best-fit parameters Ψ^*\**^ on which the reconstructed SCP geometry is closest to the image.

To numerically solve the non-linear optimization problem, we use the **fmincon** function in the *MATLAB* package, where the default algorithm is the interior-point method. To ensure a good convergence of the optimization, we set the initial guesses of parameters based on their geometrical meanings. First, the parameter ***q***_*α*_ roughly represents the cell position, so we use the cell center from the image as the initial guess. Second, the parameter *θ*_*α*_ roughly represents the cell size, so we use the cube root of the cell volume from the image as the initial guess. Third, the parameter *p*_*α*_ roughly determines the convexity of cell membranes, so we set the initial guesses randomly but following the order of all cell convexities. Additionally, we fix the parameters of the ‘background cell’ to 0 and fix the mean value of *p*_*α*_ to 1, so that the MSD has only one global minimum. The *MATLAB* code is available at https://github.com/siqiliuNU/ForceInferrence.

#### B. Verification on synthetic data

In order to verify the numerical scheme, we generate in-silico three-dimensional image data and investigate the performance of the scheme. First, we randomly set up the GWV parameters Ψ_0_ of 100 cells in a certain space region. Second, we set the size and the resolution in three dimensions, generating an image with 500*×* 500*×* 500 pixels. Third, we label each pixel with a cell index according to the GWV diagram with parameters Ψ_0_. Now a synthetic image with the segmentation of cells is constructed. In order to simulate the noise in real image data, for each membrane pixel, we arbitrarily include a Gaussian noise to the coordinate. We control the variation of the Gaussian noise from 0% up to 20%.

Then we apply the numerical scheme to the synthetic three-dimensional image using the noisy pixel coordinates. The MSD function is able to converge to a minimum at Ψ^*\**^ when using the initial guesses from our strategy. Comparing the fitted parameter Ψ^*\**^ to the ground truth Ψ_0_, we compute the accuracy of numerical scheme as

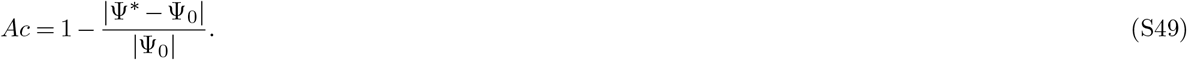

Across different levels of Gaussian noise, the inferred Ψ^*\**^ can recover the ground truth Ψ_0_ with more than 99% accuracy. This indicates that MSD minimization is a precise numerical scheme. Dive into the three classes of parameters, we find that the fitted *q*_*α*_ and *θ*_*α*_ have high accuracy, while the inference of parameter *p*_*α*_ has relatively lower precision.

#### C. Sensitivity analysis

In order to learn the robustness of the numerical scheme, we process the sensitivity analysis on the MSD function and compute the dependency of fitted parameters on the pixel noise. At the minimum of the MSD function, the derivative of *E*(Ψ, *r*) ought to be zero:

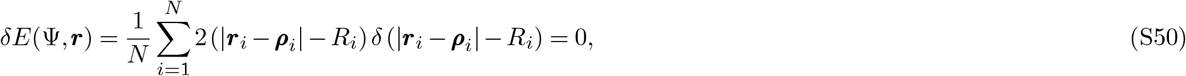

So for any pixel coordinate *r*_*i*_, there is *δ* (|*r*_*i*_ − *ρ*_*i*_(Ψ)|− *R*_*i*_(Ψ)) = 0. This deviation depends on both input *δ****r*** and the output *δ*Ψ, expressed as

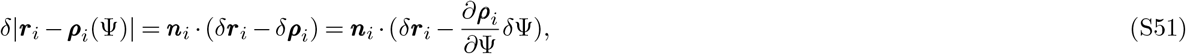

where ***n***_*i*_ = (***r***_*i*_ *−* ***ρ***_*i*_)*/*|***r***_*i*_ *−* ***ρ***_*i*_| is the normal direction of the sphere. Therefore, the deviation at ***r***_*i*_ is

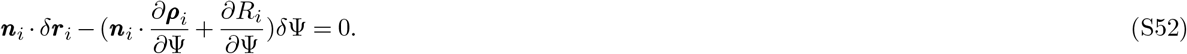

Writing all these equations together, we get a linear approximation at the minimum *Kδ****r*** + *Mδ*Ψ = 0. The vector *δ****r*** = [*δr*_1*x*_, *δr*_1*y*_, *δr*_1*z*_, …, *δr*_*ix*_, *δr*_*iy*_, *δr*_*iz*_, …]^*T*^ is the deviation of pixel coordinates. And the elements of matrix *K* are

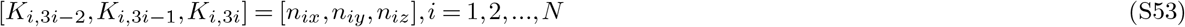

and *K*_*i,j*_ = 0 otherwise. Similarly, *δ*Ψ = [*δp*_1_, *δq*_1*x*_, *δq*_1*y*_, *δq*_1*z*_, *δθ*_1_, …, *δp*_*α*_, *δq*_*αx*_, *δq*_*αy*_, *δq*_*αz*_, *δθ*_*α*_, …]^*T*^ is the uncertainty of GWV parameters. Thus the elements of the matrix *M* is

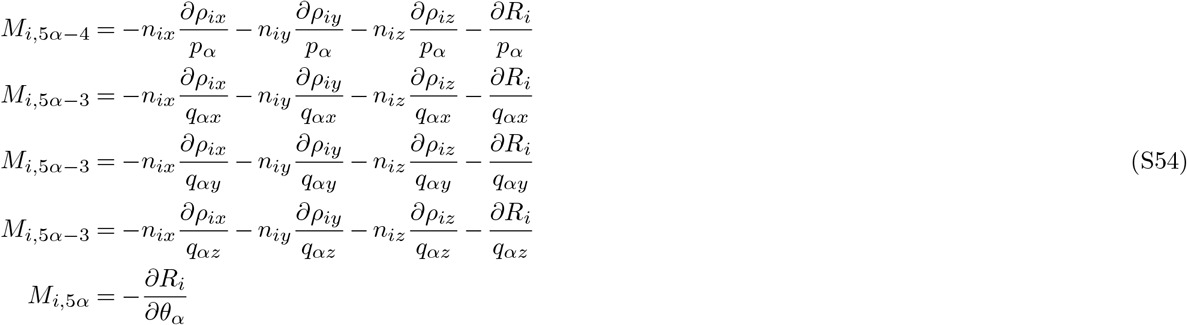

where *i* = 1, 2,…*N* ; *α* = 1, 2, …, *N*_*C*_.Therefore, the dependency between *δr* and *δ*Ψ is given by 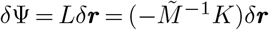, where 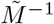 is the pseudoinverse matrix of *M*.

We use the synthetic image data to compute the dependency matrix *L*. Then we compute all the eigenvalues *λ*_*j*_ of matrix *L*. The result shows that all the eigenvalues *λ*_*j*_ are smaller than 1. This indicates that the GWV parameters Ψ are not sensitive to the pixel coordinates *r*. Therefore, the numerical scheme is a robust method to recover the geometric parameters.

**Figure S4.**
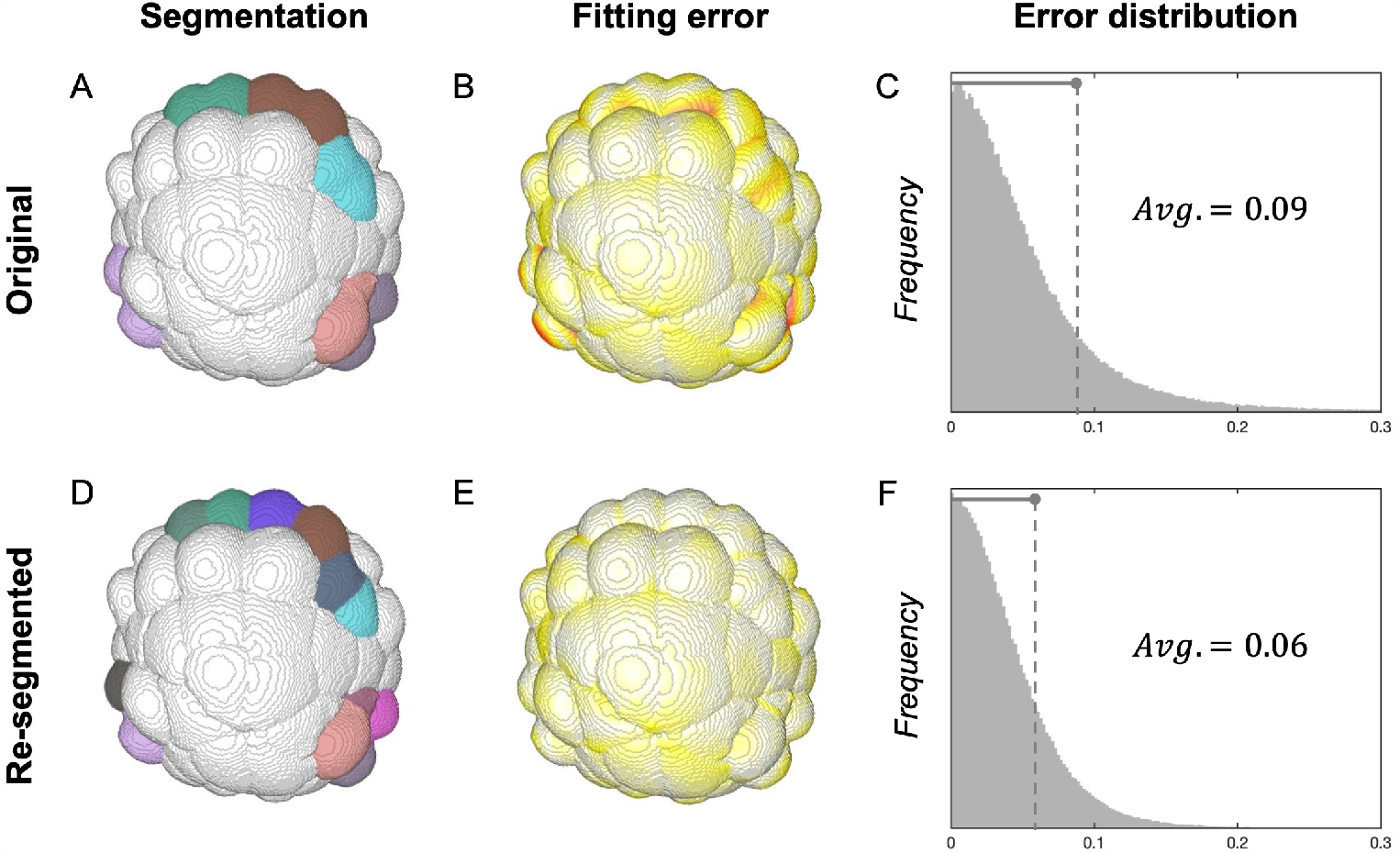
Image processing and geometry reconstruction. (A) The original segmentation of cells in a three-dimensional image during the cell-division phase. (D) The three-dimensional image with re-segmentation on dividing cells. (B)(E) The spatial distribution of fitting errors on the original image and the re-segmented image. (C)(F) The overall error distributions of the original image and the re-segmented image.

### Part IV: Force inference results on Ascidian gastrulation videos

#### A. Image processing and geometry reconstruction

We apply the force inference method to the three-dimensional image data from the study by Guignard et al [25]. We mainly focus on the embryo *ASTEC-Pm1* from the early 64-cell stage (T1) to the late 112-cell stage (T34). The images are in the same form as our synthetic images, i.e. they are three-dimensional pixel matrices where each pixel is labeled by a cell index. The image resolution and the cell numbers are also on a similar scale. Therefore, we could build the list of the membrane pixel coordinates and the corresponding cell indices, and then directly use the numerical scheme to recover the geometric parameters Ψ. At the same time, we compute the deviation *E*_*i*_ of each membrane pixel and the overall MSD minimum value for error analysis. We normalize the errors by the average size of all cells and present the distributions across space and time.

For the images during the cell-division phases, the error is relatively higher than at other time points. The high fit errors come from the dumbbell-like shapes of the cell which is about to divide. For example, as shown in Figure S4A, during the cell-division phase between 64-to 76-cell stages, the membrane shape of colored cells is far from spheres. Therefore these membranes have large fit errors (Figure S4B), thus the error distribution is wide (Figure S4C). In this case, we re-segment these dividing cells by comparing them to the image at the subsequent time point (Figure S4D). Then the fit errors are reduced to the normal level, as shown in Figure S4E and S4F.

#### B. Construction of the mechanical atlas

Once get the geometric parameters Ψ, we can analytically infer the force values on each cell, each membrane, and each edge. However, the solution of force values includes three undetermined global constants - (*a, b, c*). While the first two constants can be generically set to *a* = 0 and *b* = 1, the different choices of the third constant *c* will result in different mechanical patterns. We first investigate the upper boundary of the third constant *c*_*max*_, since it could vary across all time points. As shown in Figure S5A, the upper bound *c*_*max*_ fluctuates between the value of 0.1*−*0.2 over time. Therefore, we set the constant *c* = 0.05 and fix it over time to present the mediant case between the surface-tension-dominated pattern and the line-tension-dominated pattern.

Based on the inferred force values, we construct the mechanical atlas and visualize it using the *jet* colormap. The three-dimensional view is presented via the image analysis software *FIJI*. For each stage, we take one certain image as a representation of the mechanical pattern - T2 for the 64-cell stage, T16 for the 76-cell stage, and T34 for the 112-cell stage. One of the main patterns is the correlation between cellular pressure and cellular contractile stresses. In Figure S5B, we plot the isotropic part of the cellular contractile stress tensor (*σ*^(*c*)^) versus the cellular pressure, for all cells across all time points. The correlation coefficient is 0.89, quantitatively showing the correspondence between different mechanical inputs. We also construct the mechanical atlas for another two embryos - *ASTEC-Pm3* and *ASTEC-Pm5*. By comparing the pressure atlas in Figure S5C, we can make the qualitative statement that the mechanical patterns are reproducible across all three embryos.

**Figure S5.**
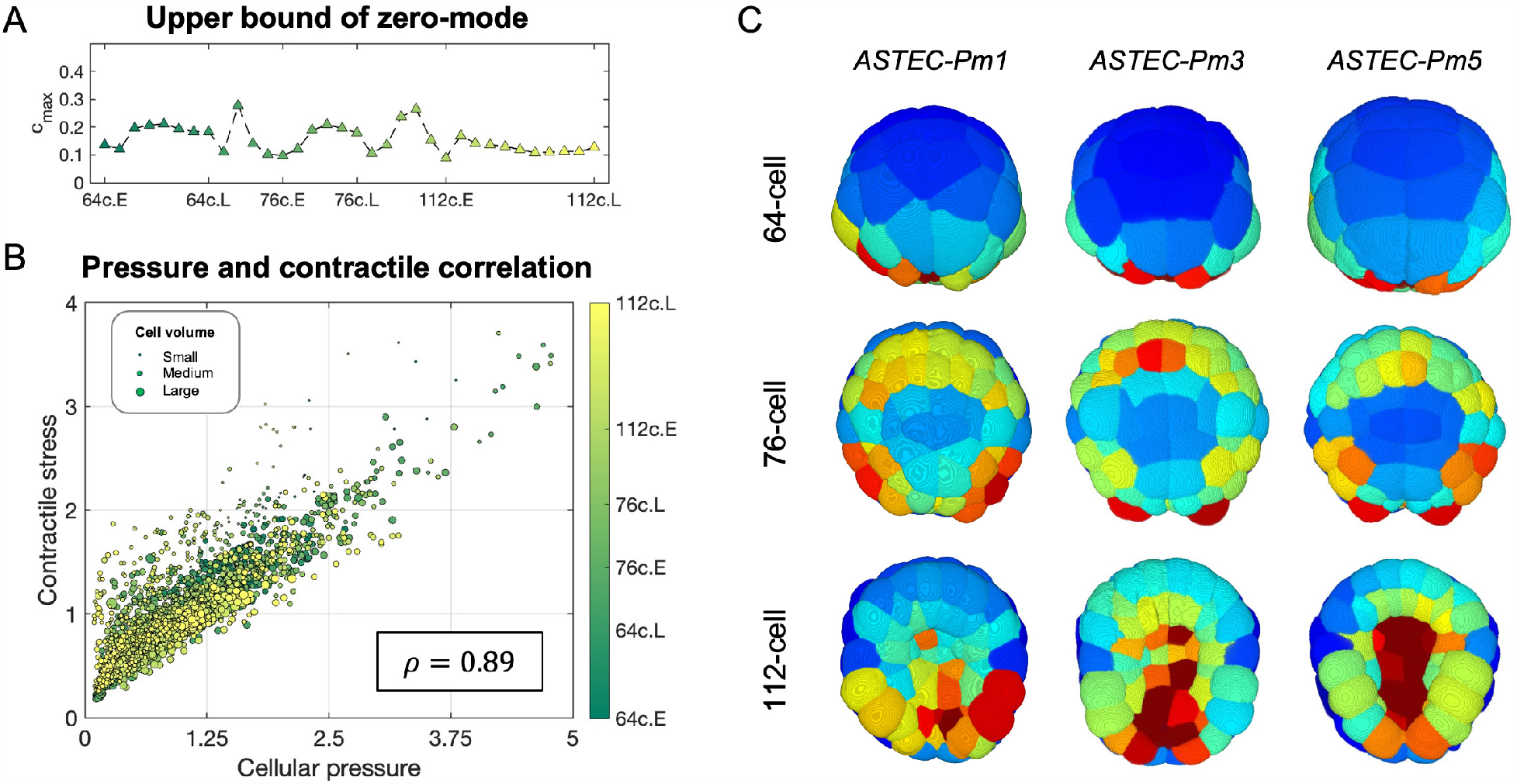
Construction of mechanical atlas. (A) The upper bound of the third mechanical zero-mode across all time points from the 64-to 112-cell stage. (B) The correlation between cellular pressures and cellular contractile stresses over all time points. (C) The reproducibility of inferred mechanical atlas across three embryos.

#### C. Quantifications of mechanical patterns

We then quantify the mechanical patterns concerning cell fates and cell positions. The cell information can be found at https://morphonet.org/. First, we quantify the symmetric and asymmetric patterns in the embryo. In Figure S6A, the comparison between the left and right corresponding pressures shows high correlation coefficients (*>* 0.9) across three stages, representing the high symmetry with respect to the left-right axis. In Figure S6B, comparing the distributions of cellular pressures in two halves along the anterior-posterior axis shows the asymmetric mechanical patterns in different stages. The posterior half has higher cellular pressures, but the difference is dimed till the 112-cell stage. In Figure S6C, comparing the distributions of cellular pressures in two halves along the animal-vegetal axis shows asymmetric mechanical patterns as well. The vegetal half has higher cellular pressures, and the difference is increased from the 76-to 112-cell stage.

Considering the detailed correlations between cellular pressures and surface or line tensions in different layers, we plot the comparisons in Figure S6D-F and Figure S6G-I. In Figure S6D, the comparison shows the high correlation between cellular pressures and apical surface tensions across all three stages. In Figure S6E, the comparison also shows the correlation between lateral surface tensions and the mean pressure in the two corresponding cells. In Figure S6F, the comparison shows the correlation between basal surface tensions and the corresponding cellular pressures on the vegetal side.

Each apical junction corresponds to two cells. In Figure S6G, we see the correlations between the apical line tensions and the mean pressure of the two corresponding cells. For lateral or basal edge junctions, each line tension corresponds to three cellular pressures. Figure S6H shows the correlations between the line tensions and the second-highest corresponding cellular pressure. This correlation is more clear when we dive into the line tensions of basal junctions in Figure S6I. A basal junction can be formed by 2 animal cells + 1 vegetal cell, or formed by 1 animal cell + 2 vegetal cells. The line tensions in the former class have a similar pattern to the pressures on the animal side, while the line tensions in the latter class have a similar pattern to the pressures on the vegetal side. In summary, Figure S6D-I presents the related patterns between three different mechanical inputs.

**Figure S6.**
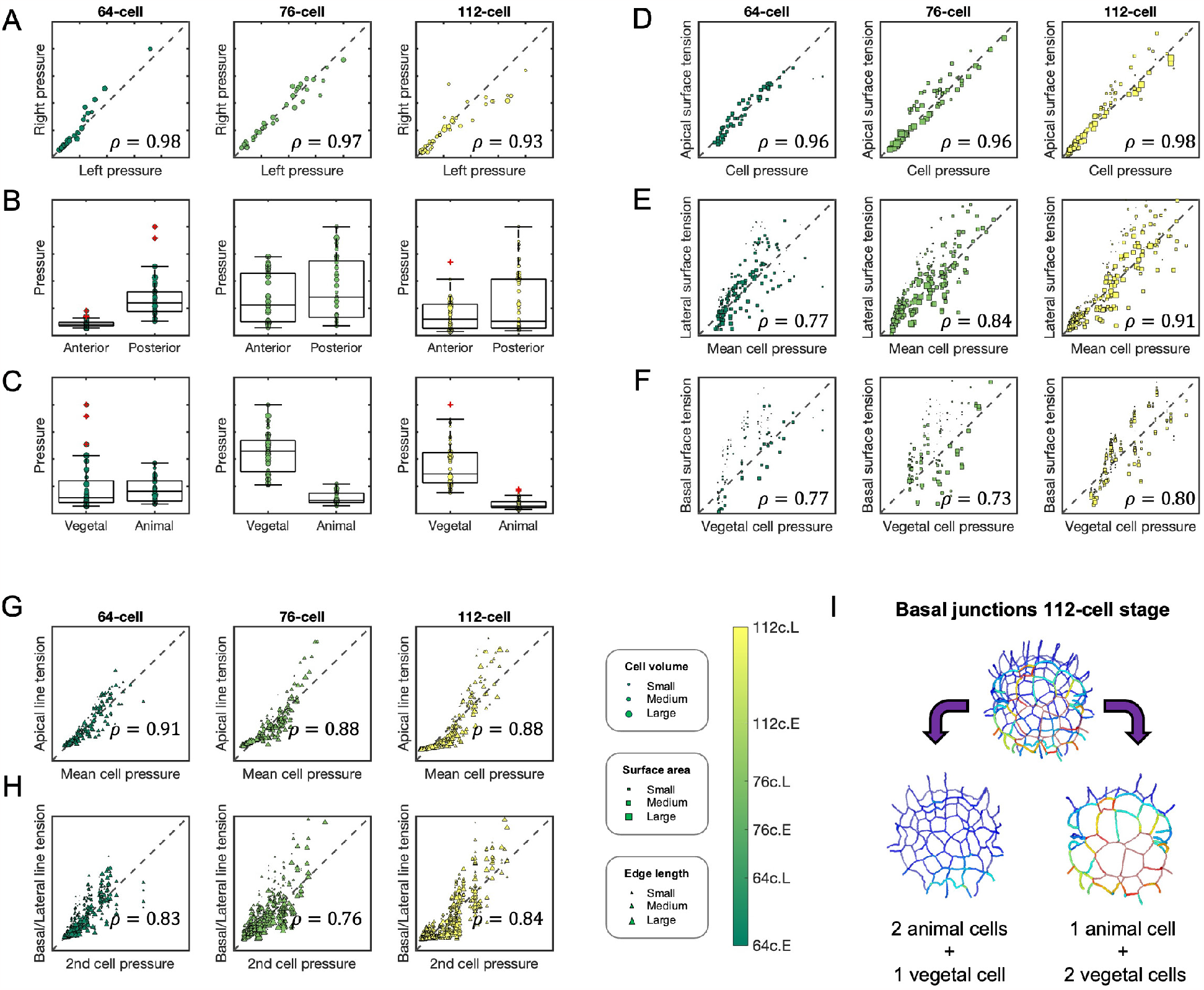
Quantifications of mechanical patterns. (A)(B)(C) The symmetry and asymmetry quantifications of cellular pressures with respect to LR, AP, and AV axes, across three stages. (D)(E)(F) The correlations between surface tensions and cellular pressures in three classes of membranes, across three stages. (G)(H) The correlations between line tensions and cellular pressures in two classes of junctions, across three stages. (I) The two sub-classes of basal junctions at the 112-cell stage.

#### D. Mechanical patterns of von Mises stress

As introduced above, the von Mises stress captures the anisotropic component of the cellular stress tensor. We heatmap the variation of *σ*_*V M*_ on the lineage diagram of the embryo in Figure S7B, and present its spatial patterns from an animal and vegetal view at the 112-cell stage in Figure S7A. The pattern of the von Mises stress in Figure S7B is manifestly distinct from either of the patterns seen in pressures or contractile stresses. This is naturally expected since the pressure (hydrostatic) and deviatoric contributions to overall stress are independent. Contrasting previously observed patterns, ectodermal cells experience shear stress. To gain a more physical sense of the pattern we visualize the von Mises stress on the embryo in Figure S7A. One can clearly observe patterns, including the high levels of von Mises stress in the posterior mesodermal lineage.

**Figure S7.**
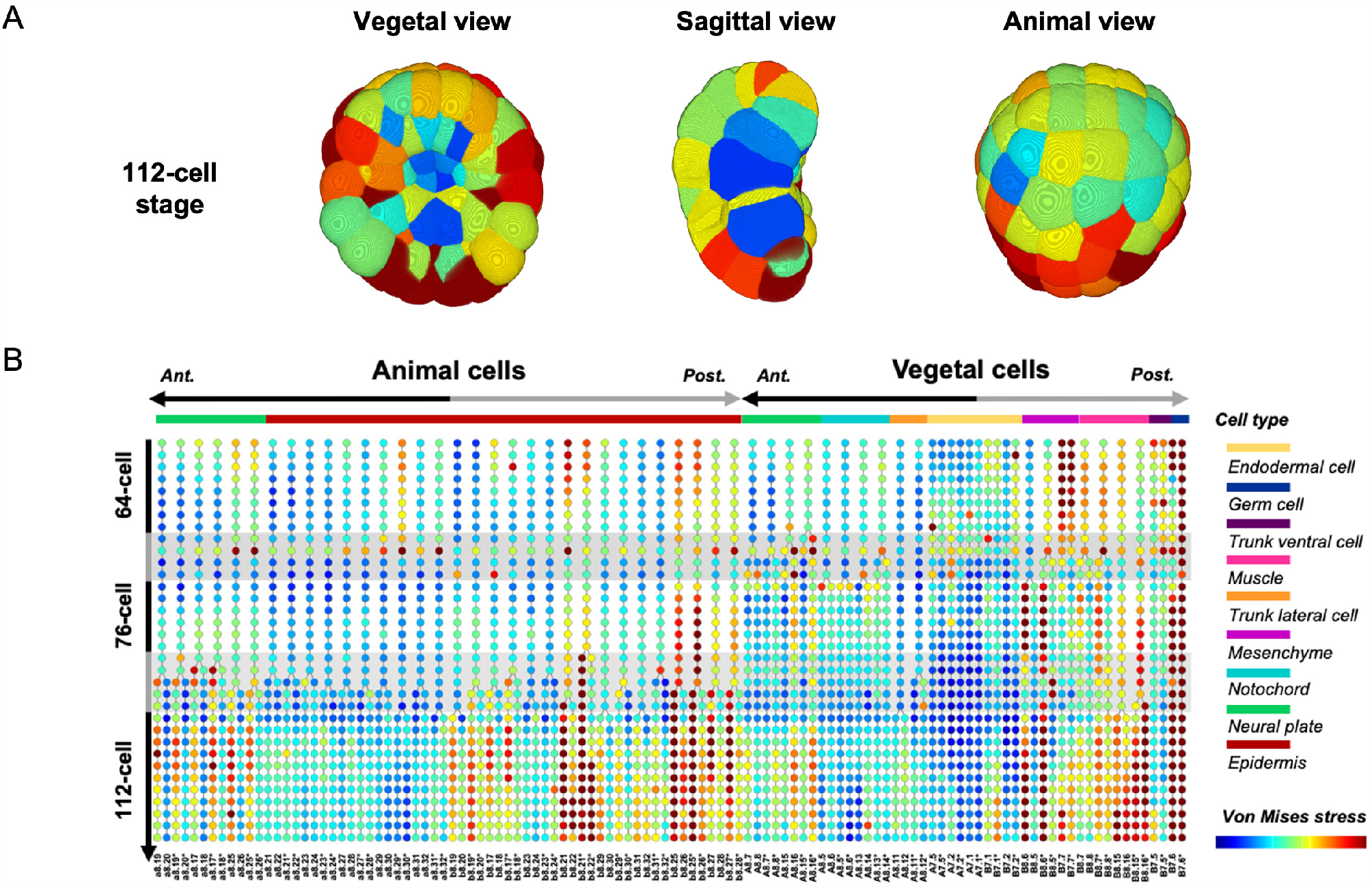
Mechanical patterns of von Mises stress. (A) The spatial pattern of von Mises stress at the 112-cell stage, in vegetal, sagittal, and animal views. (B) The lineage diagram of von Mises stress in the embryo *ASTEC-Pm1*.

